# Tissue-wide Genetic and Cellular Landscape Shapes the Execution of Sequential PRC2 Functions in Neural Stem Cell Lineage Progression

**DOI:** 10.1101/2022.04.04.487003

**Authors:** Nicole Amberg, Florian M. Pauler, Carmen Streicher, Simon Hippenmeyer

## Abstract

The generation of a correctly-sized cerebral cortex with all-embracing neuronal and glial cell-type diversity critically depends on faithful radial glial progenitor (RGP) cell proliferation/differentiation programs. Temporal RGP lineage progression is regulated by Polycomb Repressive Complex 2 (PRC2) and loss of PRC2 activity results in severe neurogenesis defects and microcephaly. How PRC2-dependent gene expression instructs RGP lineage progression is unknown. Here we utilize Mosaic Analysis with Double Markers (MADM)-based single cell technology and demonstrate that PRC2 is not cell-autonomously required in neurogenic RGPs but rather acts at the global tissue-wide level. Conversely, cortical astrocyte production and maturation is cell-autonomously controlled by PRC2-dependent transcriptional regulation. We thus reveal highly distinct and sequential PRC2 functions in RGP lineage progression that are dependent on complex interplays between intrinsic and tissue-wide properties. In a broader context our results imply a critical role for the genetic and cellular niche environment in neural stem cell behavior.

## INTRODUCTION

The mammalian neocortex is composed of a rich variety of cell types and regulates critical cognitive and behavioral functions in the brain. The majority of neocortical cells include distinct excitatory and inhibitory neuronal classes and glial cell types (Tasic et al., 2018). During cortical development excitatory projection neurons and glial cells are generated from radial glial progenitors (RGPs) in a temporal sequential manner (Beattie and Hippenmeyer, 2017; Dwyer et al., 2016; Oberst et al., 2019; Smart, 1973; Taverna et al., 2014). RGP lineage progression in the developing embryonic cortex occurs in three major consecutive stages: 1) symmetric expansion of RGP pool; 2) asymmetric neurogenic phase; and 3) initiation of glial cell production (Kriegstein and Alvarez-Buylla, 2009; Taverna et al., 2014). RGPs also generate a variety of distinct postnatal lineages (besides glial cells) including ependymal cells or postnatal neural stem cells (Gallo and Deneen, 2014). Recent quantitative single RGP cell lineage tracing by using Mosaic Analysis with Double Markers (MADM) revealed that overall RGP proliferation behavior is stereotyped and predictable to a large extent (Gao et al., 2014; Llorca et al., 2019; Shen et al., 2021; Zhang et al., 2020). In other words, RGPs undergo a defined series of symmetric proliferative divisions before switching to asymmetric neurogenic division. Once in asymmetric division mode, individual neurogenic RGPs generate a unitary output of ∼8-9 neurons before transiting to glia production with about one in six RGPs proceeding to generate glial cells (Beattie and Hippenmeyer, 2017; Lin et al., 2021). Although the above inaugural quantitative framework has been established at single cell level, the precise cellular and molecular mechanisms regulating the orderly RGP lineage progression are not well understood.

At the individual cell level, RGPs appear to undergo a series of molecular identity and/or transcriptional state changes correlating with lineage progression and the generation of correctly specified progeny (Di Bella et al., 2021; La Manno et al., 2021; Telley et al., 2019). In general, the transcriptional landscape in stem cell lineage progression is tightly regulated by epigenetic mechanisms (Amberg et al., 2019) including PRC2-mediated posttranslational chromatin modifications. PRC2 consists of three core subunits that are essential for proper catalytic activity and transcriptional repression *in vivo*: embryonic ectoderm development (EED), enhancer of zeste 2 (EZH2) or its homolog EZH1, and suppressor of zeste 12 (SUZ12) (Di Croce and Helin, 2013). Both, EZH2 and EZH1, contain a conserved SET domain capable of catalyzing the mono-, di-, and tri-methylation of lysine 27 of histone H3 (H3K27) (Di Croce and Helin, 2013).

H3K27me3 is recognized by PRC1, which catalyzes the mono-ubiquitinylation of lysine 119 of histone H2A (H2AK119ub) and thereby ultimately induces transcriptional silencing (Bracken and Helin, 2009; de Napoles et al., 2004). Ablation of any one of the PRC2 core components *Eed*, *Ezh2* or *Suz12* results in complete disruption of the complex and subsequent loss of H3K27me3 (Montgomery et al., 2005; Pasini et al., 2004), disrupting embryonic development (Leeb and Wutz, 2007). In effect full gene knock-out of either, *Eed*, *Ezh2* or *Suz12* results in early embryonic lethality (Faust et al., 1998; Niswander et al., 1988; Pasini et al., 2004).

PRC2 activity is critical for the orchestration of cortical projection neuron and glial cell generation (Hahn et al., 2013; Hirabayashi et al., 2009; Pereira et al., 2010; Telley et al., 2019; Zhao et al., 2015). Conditional loss (cKO) of PRC2 function by ablation of *Ezh2* (Pereira et al., 2010) or *Eed* (Telley et al., 2019) in *Emx1^+^* RGPs results in dramatic thinning of the developing cortical wall with concomitant microcephaly. In correlation with the strong microcephaly observed in *Eed* cKO, RGPs seem to show accelerated temporal progression with shortened neurogenic period (Telley et al., 2019). At later developmental stages, absence of PRC2 in RGPs was shown to affect the neurogenic-to-gliogenic switch (Hirabayashi et al., 2009; Pereira et al., 2010). However, the exact function of PRC2 in astrocyte generation is still under debate because Hirabayashi et al. (Hirabayashi et al., 2009) observed delayed gliogenesis in *Ezh2* cKO whereas Pereira et al. (Pereira et al., 2010) reported precocious onset of gliogenesis. The discrepancy of the above findings may reflect a putative dual role for PRC2 in regulating major transitions in RGPs at both neurogenic and gliogenic competence levels, albeit concrete evidence for such mode of PRC2 action is currently lacking (Hirabayashi et al., 2009; Pereira et al., 2010; Telley et al., 2019). In summary, based on whole tissue ablation, PRC2 has been proposed to exert several critical roles in cortical RGPs. How PRC2 regulates RGP proliferation behavior at the individual cell level in distinct stages along their lineage progression remains unknown.

Due to its cell-intrinsic mode of action, PRC2 activity is generally thought to be a cell-autonomous one. Yet, individual RGPs *in vivo* are embedded within the neocortical ventricular stem cell niche and thus operate in a complex cellular environment. Whether global tissue-wide properties shape the cell-autonomous PRC2 requirement or function in individual RGPs is currently unclear. Previous studies mainly utilized experimental paradigms with global and/or whole tissue-wide loss of PRC2 function, and analysis of RGP lineage progression lacked single cell resolution. To overcome this limitation, in this study we used Mosaic Analysis with Double Markers (MADM) technology in order to analyze the functional requirement of PRC2 at individual RGP cell level. MADM enables the generation of sparse mutant clones (Beattie et al., 2020; Contreras et al., 2021; Zong et al., 2005) and thus the study of true cell-autonomous PRC2 requirement. We contrasted the sparse PRC2 elimination with whole tissue ablation paradigm and genetically dissected the interplay of PRC2 cell-autonomous and global tissue requirement. Contrary to its predicted essential function in embryonic neurogenic RGPs we could show that PRC2 is not cell-autonomously required in the control of neurogenesis but rather plays a critical role at the global tissue-wide level. In contrast, at postnatal stages PRC2 was cell-autonomously required for astrocyte production and maturation. Altogether, our data revealed distinct sequential, cell-autonomous and global tissue-wide functional PRC2 requirements in RGP lineage progression during cortical development.

## RESULTS

### Global but not sparse KO of PRC2 results in diminished cortical projection neuron production and microcephaly

In order to obtain insights of PRC2 function in RGPs with single cell resolution we combined a floxed allele of the core subunit *Eed* [*Eed*-flox; (Yu et al., 2009)] with MADM cassettes located on chr.7 (MADM-7) (Hippenmeyer et al., 2013) (Figure S1). In a first experiment we conditionally removed *Eed* in a global tissue-wide manner and generated *Eed* cKO mice, with concomitant sparse MADM labeling, in combination with *Emx1*-Cre (Gorski et al., 2002) (*MADM-7^GT,Eed/TG,Eed^;Emx1^Cre/+^* abbreviated cKO-*Eed*-MADM). As expected and previously reported (Pereira et al., 2010; Telley et al., 2019) cKO-*Eed*-MADM mice at postnatal day (P) 21 showed a strong decrease of neocortical thickness when compared to control-MADM (*MADM-7^GT/TG^;Emx1^Cre/+^*) animals (Figures 1A-B, 1D-1G, 1J and 1L). In both control-MADM (all cells *Eed^+/+^*) and cKO-*Eed*-MADM (all cells *Eed^-/-^*) the green (GFP^+^) to red (tdT^+^) (g/r) ratio was ∼1 reflective of the identical genotypes in GFP^+^ and tdT^+^ MADM-labeled cells (Figures 1K and 1M). The dramatic thinning of the neocortex in cKO-*Eed*-MADM reflects a decrease of neuron production, presumably due to accelerated lineage progression of neurogenic RGPs (Telley et al., 2019). Thus, assuming a purely cell-autonomous function of PRC2 in proliferating RGPs, we predicted a dramatic loss of *Eed* mutant cells in comparison to wild-type cells when analyzed in a sparse genetic mosaic as provided by the MADM system (Figure 1C). To test the above, we generated mosaic *Eed*-MADM (*MADM-7^GT/TG,Eed^;Emx1^Cre/+^*) mice with sparse green GFP^+^ homozygous *Eed^-/-^* mutant cells, yellow (GFP^+^/tdT^+^) heterozygous *Eed^+/-^*, and red (*tdT^+^*) *Eed^+/+^* wild-type cells in an otherwise unlabeled heterozygous background (Figures 1C, 1H, 1I and S1D-S1E) (Amberg and Hippenmeyer, 2021). Contrary to our expectation, we found no reduction of green *Eed^-/-^* mutant when compared to red *Eed^+/+^* control neurons and thus a g/r ratio of ∼1 at P21 (Figure 1O). To validate the specificity of PRC2 activity elimination in green *Eed^-/-^* mutant but not red *Eed^+/+^* control cells in mosaic *Eed*-MADM mice, we stained for the PRC2-catalyzed histone mark H3K27me3 in E12.5 cortex (Figure S2). H3K27me3 signal and thus PRC2 activity was abolished in individual green *Eed^-/-^* but not red *Eed^+/+^* cells in mosaic *Eed*-MADM brains (Figure S2 C-S2F). In contrast and as expected, H3K27me3 was completely eliminated in all cells in cKO-*Eed*-MADM (Figure S2 G-S2J).

**Figure 1.**
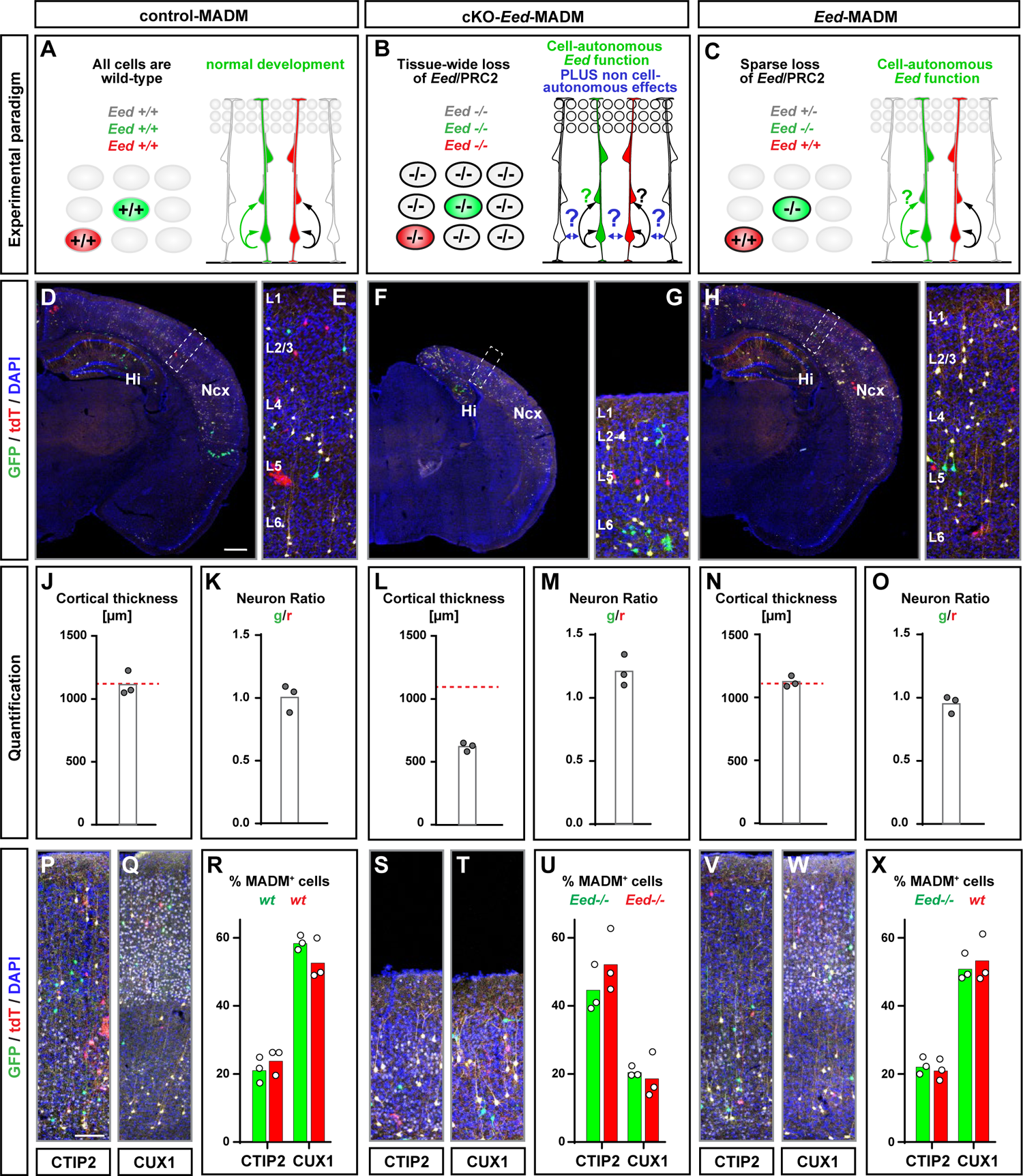
*Eed*/PRC2 is not cell-autonomously required in cortical neurogenesis. See also Figures S1 and S2. **(A-C)** Schematic overview of experimental paradigm and genotype of MADM-labeled cells in (A) wild-type MADM (control-MADM); (B) conditional knockout of *Eed* in all cortical projection neurons (cKO-*Eed*-MADM); and (C) sparse genetic mosaic with *Eed* mutant cells labeled in green color (*Eed*-MADM). **(D-I)** Overview of MADM-labeling pattern in somatosensory cortex in (D) control-MADM; (F) cKO-*Eed*-MADM; and (H) *Eed*-MADM mice at P21. (E, G, I) depict higher resolution images of boxed areas in (D), (F) and (H) with indication of cortical layers. **(J-O)** Quantification of cortical thickness and green-to-red (g/r) neuron ratio, respectively, in (J-K) control-MADM; (L-M) cKO-*Eed*-MADM; and (N-O) *Eed*-MADM. **(P-X)** Histological stainings of P21 brains for lower-layer marker CTIP2 and for upper-layer marker CUX1, and quantification of the percentage (%) of green and red MADM-labeled CTIP2^+^ and CUX1^+^ cells in (P-R) control-MADM; (S-U) cKO-*Eed*-MADM; and (V-X) *Eed*-MADM mice. Each individual data point represents one experimental animal. Data indicate mean ± SEM. Scale bars: 500µm in (D, F, H); 60µm in (E, G, I, P, Q, S, T, V, W).

Next we analyzed the generation of CTIP2^+^ (primarily lower layer V) relative to CUX1^+^ (upper layers IV-II) neurons. While we found very similar relative fractions of CTIP2^+^ and CUX1^+^ MADM-labeled cells in control-MADM and mosaic *Eed*-MADM (Figures 1P-1R and 1V-1X), the relative abundance of CTIP2^+^ MADM-labeled cells was significantly increased when compared to CUX1^+^ cells in cKO-*Eed*-MADM in agreement with previous studies (Telley et al., 2019) (Figures 1S-1U). We conclude that global KO of *Eed* in cKO-*Eed*-MADM lead to dramatic microcephaly due to significant reduction of upper layer IV-II relative to lower layer V projection neurons. In contrast, sparse deletion of *Eed* in genetic mosaic *Eed*-MADM paradigm did not affect projection neuron production and thus cortical size. Together, the above data suggested that *Eed* is not cell-autonomously required in cortical RGPs during neurogenic phase but rather at the global tissue-wide level.

### PRC2 function is not cell-autonomously required in cortical RGP-mediated projection neuron production

While the above analysis at the population level already provided an indication about cell-autonomy of *Eed* gene function based on sparseness of the genetic mosaic, conclusive assessment of PRC2 function at the individual RGP level is only possible using clonal analysis. Thus we carried out MADM-based clonal analysis (Beattie et al., 2020) to probe cell-autonomous PRC2 function at true single RGP cell level. To determine the neurogenic potential of individual mutant *Eed^-/-^* RGPs we pursued two MADM assays in combination with tamoxifen (TM)-inducible *Emx1^CreER^* driver (Kessaris et al., 2006).

First, we analyzed RGPs in their symmetric proliferative mode (Gao et al., 2014). In the MADM context, the two daughter cells (and resulting subclones) from such symmetric proliferative RGP divisions will be labeled in different colors (red and green) and may inherit distinct genotypes (Beattie et al., 2020). We first validated the approach since MADM clones using MADM-7 in combination with *Emx1^CreER^* driver had not been much analyzed previously. We thus first generated control *MADM-7^GT/TG^;Emx1^CreER^* and injected TM at E11.5 when the majority of RGPs still divide symmetrically (Gao et al., 2014). We quantified the cell numbers in the two red and green (both wild-type) subclones at E13.5 across the developing cortical wall (Figure 2A-2C); and at E16.5, when most neurons have been generated (Figure 2G-2I). As expected, the numbers of green and red wild-type neurons at both E13.5 and E16.5 and in all analyzed zones was not significantly different (Figure 2C and 2I). Next, we generated *Eed*-MADM clones (*MADM-7^GT/TG,Eed^;Emx1^CreER^*) with similar TM injection/analysis regime as above but with the red subclone containing wild-type and the green subclone *Eed^-/-^* mutant projection neurons, respectively. At both E13.5 and E16.5 analysis time points we counted similar neuron numbers like in the above control-MADM clones. We could not detect significantly different numbers of red wild-type compared to green *Eed^-/-^* mutant neurons (Figure 2D-2L). Thus, the proliferation potential of mutant *Eed^-/-^* and wild-type subclones emerging from a single symmetrically dividing RGP appeared identical.

**Figure 2.**
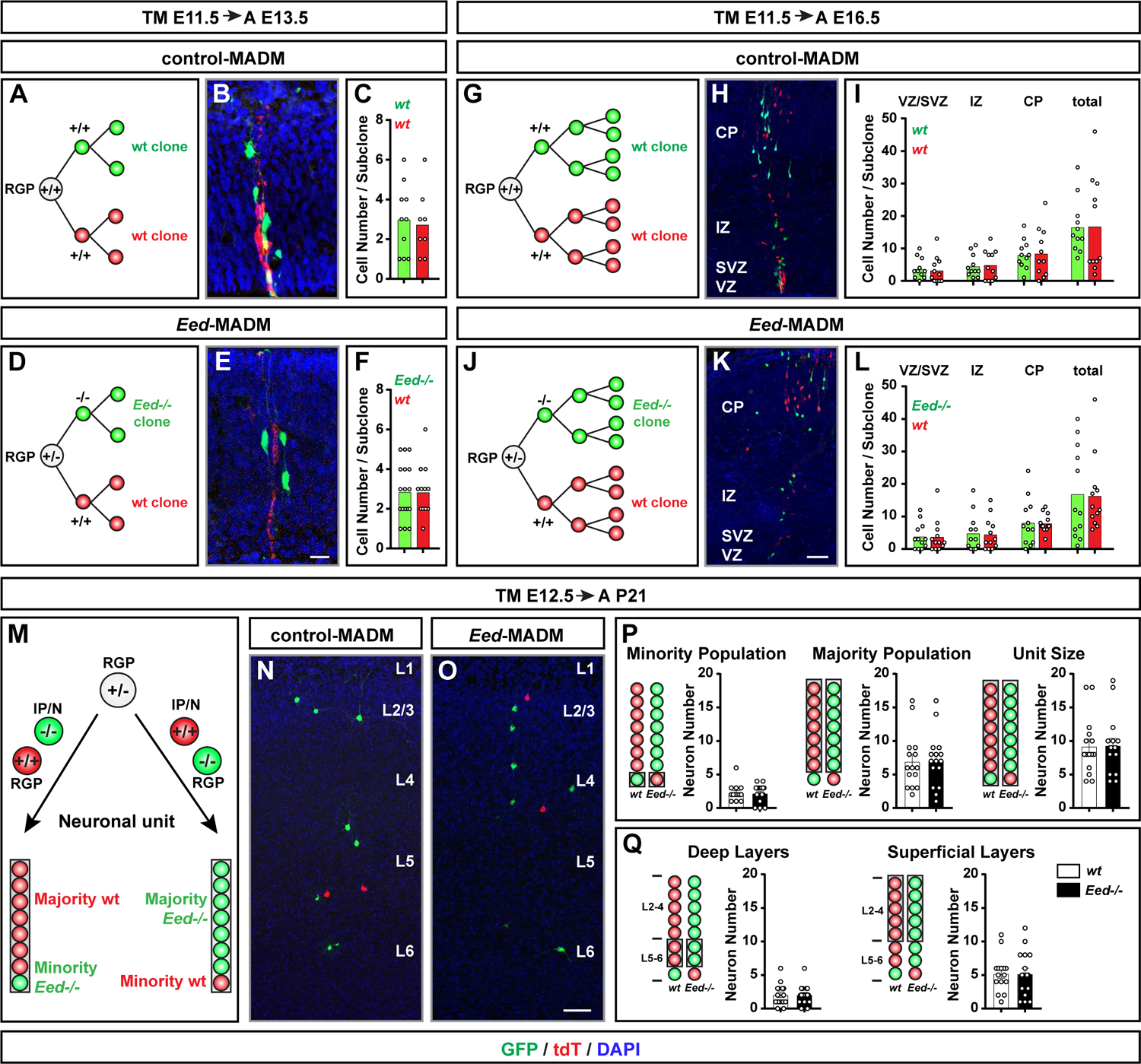
Functional analysis of *Eed*/PRC2 function during neurogenesis at single RGP clone level. **(A-F)** Experimental setup for embryonic MADM clones induced at E11.5 and analyzed at E13.5 with representative images and corresponding quantifications in (A-C) control-MADM; and (D-F) *Eed*-MADM mice. **(G-L)** Experimental setup for embryonic MADM clones induced at E11.5 and analyzed at E16.5 with representative images and corresponding quantifications in (G-I) control-MADM; and (J-L) *Eed*-MADM mice. **(M)** Experimental setup for MADM clones induced at E12.5 and analyzed at P21 to determine the neuronal unit size. Upon a MADM event in asymmetrically dividing RGPs, one daughter cell will become an intermediate progenitor or neuron (IP/N) and thus constitute the minority population of the clone, while the other daughter cell will remain RGP identity and produce the majority population of the clone. **(N-O)** Representative images of clones in (N) control-MADM; and (O) *Eed*-MADM mice. **(P)** Quantification of the average neuron number within the minority population (left panel), the majority population (middle panel) and the total unit size (right panel) emerging from wild-type and *Eed^-/-^* RGPs. **(Q)** Quantification of the average number of deep layer neurons (left panel) and superficial layer neurons (right panel) in wild-type and *Eed^-/-^* MADM clones. Each individual data point represents one MADM clone. Data indicate mean ± SEM. Scale bars: 20µm in (B, E); 50µm in (H, K, N, O).

In the second assay we quantified absolute neuron output of individual RGPs once they switched to asymmetric neurogenic proliferation mode (Gao et al., 2014). Previous MADM analysis, using MADM-11, has demonstrated unitary output of ∼8-9 neurons from a single RGP (Beattie et al., 2017; Gao et al., 2014; Llorca et al., 2019). We injected TM at E12.5 and analyzed MADM clone composition at P21 (Figure 2M-2O). We analyzed total unit size, majority population (larger subclone) and minority population (smaller subclone) (Figure 2P), and distribution of MADM-labeled neurons in deep (VI-V) and superficial (IV-II) cortical layers (Figure 2Q). In all quantifications we could not detect a significant difference in the numbers of *Eed^-/-^* mutant when compared to *Eed^+/+^* control neurons, and the total unit size was ∼8-9 neurons for each genotype. Based on the above results from single MADM clone analysis we conclude that PRC2 function is not cell-autonomously required for cortical RGP-mediated neuron production.

### Distinct tissue-wide genetic environment in mosaic and global KO triggers differential ectopic gene expression in *Eed^-/-^* mutant cells

Although *Eed^-/-^* cells in *Eed*-MADM and cKO-*Eed*-MADM paradigms had identical genotypes they existed in distinct genetic and cellular environments. While in *Eed*-MADM the genetic background predominantly was *Eed*^+/-^ heterozygous with phenotypically regular sized cortex (Figure 1N), the vast genetic landscape in cKO-*Eed*-MADM is *Eed*^-/-^ with massively diminished neuron numbers and much smaller overall cortex size (Figure 1L). Given the dramatic phenotypic difference in individual MADM-labeled *Eed^-/-^* mutant cells depending on their genetic and cellular environment we sought to identify molecular correlates hinting at global tissue-wide deregulation of gene expression due to PRC2 loss of function. We utilized an established FACS-based approach (Laukoter et al., 2020a; Laukoter et al., 2020c) to isolate MADM-labeled *Eed*^+/+^, *Eed*^+/-^ and *Eed*^-/-^ mutant cells in a time course at E12.5 (onset of neurogenesis), E13.5 (early neurogenesis), E16.5 (end of neurogenesis) and P0 (start of gliogenesis) (Figures 3A-3D). Next we performed small sample (up to 400 cells) Smart-Seq2-based bulk RNA sequencing followed by transcriptome analysis. We confirmed that *Eed* expression was efficiently depleted in *Eed^-/-^* mutant cells in *Eed*-MADM and cKO-*Eed*-MADM at all time points (Figure S3 A-B). Principle component analysis (PCA) revealed that individual samples clustered according to age and genetic paradigm (Figure S3 C). Interestingly, samples of green cells from control-MADM (*Eed*^+/+^) and *Eed*-MADM (*Eed*^-/-^) were found in same clusters at all time points while samples of green cells from cKO-*Eed*-MADM (*Eed*^-/-^) formed distinct clusters from E13.5 onwards (Figure S3 C). The PCA analysis indicated that *Eed*^-/-^ mutant cells from *Eed*-MADM showed a more similar transcriptome to wild-type but a distinct transcriptome when compared to *Eed*^-/-^ cells from cKO-*Eed*-MADM. Thus we aimed to directly test whether variable global gene expression in mutant *Eed*^-/-^ cells correlateed with the genetic/cellular environment. We identified differentially expressed genes (DEGs) of *Eed^-/-^* cells between *Eed*-MADM and cKO-*Eed*-MADM. A relatively low number of DEGs was found at E12.5 (42) and E13.5 (160) but increasing numbers of DEGs were present at E16.5 (1980) and P0 (4217). DEGs showed temporal developmental specificity with limited overlap between the different time points (Figure S3 D).

**Figure 3.**
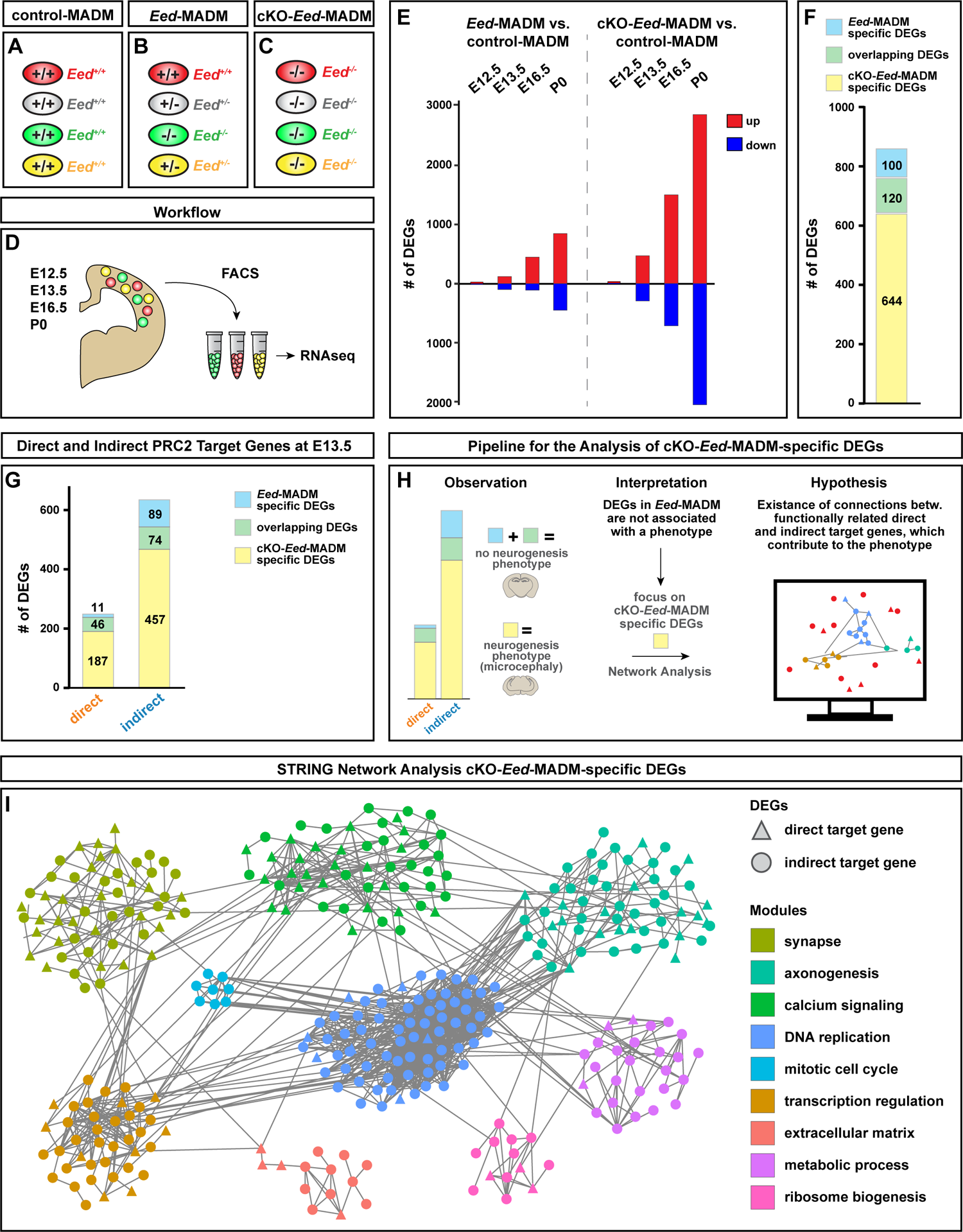
Deregulated genes specific to cKO-*Eed*-MADM are functionally connected and converge on cell cycle regulatory modules. See also Figure S3 and S4. **(A-C)** Schematic overview of experimental MADM paradigms and genotype of differentially labeled cells in (A) control-MADM; (B) *Eed*-MADM; and (C) cKO-*Eed*-MADM. **(D)** Schematic overview of workflow for RNA-seq experiments at E12.5, E13.5, E16.5 and P0. **(E)** Number of DEGs (padj < 0.1, DESeq2) upon comparison of *Eed^-/-^* mutant (*Eed*-MAD, cKO-*Eed*-MADM) cells to *Eed^+/+^* cells in control-MADM at different developmental time points (E12.5, E13.5, E16.5 and P0). ‘Up’ refers to genes upregulated in *Eed^-/-^* mutant cells, while ‘down’ refers to genes downregulated in *Eed^-/-^* mutant cells when compared to *Eed^+/+^* cells. **(F)** Detailed analysis of E13.5 DEGs. Number of DEGs specific to *Eed^-/-^* cells from *Eed*-MADM (light blue), specific to *Eed^-/-^* cells from cKO-*Eed*-MADM (light yellow) and genes shared between both paradigms (light green). **(G)** Overlap analysis of E13.5 DEGs (same as in F) with H3K27me3 data (Albert et al., 2017). Number of direct and indirect PRC2 target genes specific to *Eed^-/-^* cells from *Eed*-MADM (light blue), specific to *Eed^-/-^* cells from cKO-*Eed*-MADM (light yellow) and genes shared between both paradigms (light green). Note that only genes informative in both datasets have been included in the analysis. For details see text and Materials and Methods. **(H)** Schematic illustration of hypothesis-based STRING network analysis of cKO-*Eed*-MADM-specific DEGs. **(I)** STRING network showing protein-protein interactions of cKO-*Eed*-MADM-specific DEGs at E13.5. Triangles: direct PRC2 target genes, circles: indirect PRC2 target genes (same as in G), rectangle: genes not informative in (Albert et al., 2017). Note that DEGs group into functionally distinct modules as indicated by color code. For details see text and Materials and Methods.

To gain more detailed insights into gene expression changes, we compared *Eed^-/-^* cells from *Eed*-MADM and cKO-*Eed*-MADM with *Eed^+/+^* cells in control-MADM. Similar to the initial analysis we found increasing numbers of DEGs from E12.5 to P0 (Figure 3E). Consistent with the well described function of PRC2 in transcriptional repression, the majority of DEGs was comprised of upregulated genes (Figures 3E and S3D) (Bracken and Helin, 2009). Interestingly, the number of DEGs was approximately 3-4 fold higher in *Eed^-/-^* cells from cKO-*Eed*-MADM than from *Eed*-MADM at any given developmental time point (Figure 3E). Since *Eed* cKO mice were reported to display altered RGP behavior from E14.5 onwards (Telley et al., 2019), we focused further gene expression analysis on E13.5. Overlap analysis of E13.5 DEGs revealed that 644 DEGs were specifically deregulated in cKO-*Eed*-MADM, while only 100 DEGs were specific to *Eed*-MADM and 120 DEGs were shared between both paradigms (Figure 3F).

PRC2 is a direct transcriptional repressor via deposition of the H3K27me3 marks. We therefore sought to identify deregulation of genes directly caused by the lack of PRC2 (direct target genes) as well as gene expression changes caused downstream of direct targets (indirect targets). To this end, we utilized a published H3K27me3 ChIPseq dataset from purified cortical progenitor cells from E14.5 cortex (Albert et al., 2017). Genes reported to contain H3K27me3 marks and showing significant upregulation were classified as direct target genes. All other significantly deregulated genes were classified as indirect targets (Figure S4 A-B). Such analysis identified 4x more direct target genes in *Eed^-/-^* cells from cKO-*Eed*-MADM (235) than from *Eed*-MADM (58) (Figure 3G). In line with the function of PRC2 as a transcriptional repressor we found a larger number of upregulated genes when direct target genes were compared to indirect targets (Figure S4 B). DEGs identified as indirect targets were massively downregulated and much more abundant than direct targets (Figure S4 B, 108 vs 58 in *Eed*-MADM, 426 vs 235 in cKO-*Eed*-MADM). Yet, we found a greater overlap of DEGs classified as direct PRC2 target genes (47/58 or 81% in *Eed*-MADM vs 47/235 or 20% in KO-*Eed*-MADM) compared to indirect target genes (46/108 or 43% in *Eed*-MADM, 46/426 or 11% in cKO-*Eed*-MADM) (Figure 3G). Altogether, our results demonstrate that the genetic constitution of the cellular environment strongly influences the amount of transcriptionally silenced direct PRC2 target genes.

### Whole tissue KO of *Eed* lead to deregulation of strongly connected but functionally diverse gene modules

The above data suggested that cKO-*Eed*-MADM specific DEGs alone could cause cKO-*Eed*-MADM specific phenotypes for two reasons: 1) The number of specific direct target genes is over ten times higher in a mutant environment (188 in cKO-*Eed*-MADM vs. 11 in *Eed*-MADM) (Figure 3G); and 2) *Eed^-/-^* cells in cKO-*Eed*-MADM but not in *Eed*-MADM showed neurogenesis deficits. Signaling complexes perform their biological activity in the context of larger modules consisting of functionally-related and interacting proteins (Szklarczyk et al., 2021). We thus tested whether cKO-*Eed*-MADM specific DEGs contained modules that were sufficient to explain the altered RGP behavior and microcephaly phenotype. To do so, we made use of the rich information of protein-protein interactions in the STRING databases (Szklarczyk et al., 2021). We used the 644 cKO-*Eed*-MADM-specific DEGs as input to the STRING analysis (Figure 3H). Strikingly, proteins associated with our list of DEGs formed a highly significant and densely connected network (PPI enrichment p-value <10^-16^). For further analysis we focused on the largest connected subnetwork, consisting of 333 genes (see Materials and Methods). Clustering analysis of this network identified 9 modules of strongly connected genes (Figure 3I). Gene Ontology (GO) analysis of genes in each of these 9 modules revealed significant enrichment of diverse terms (Supplementary Table S1). The top 3 enriched GO terms were used to label each module (Figures 3I and S4C). Importantly, two modules held the potential to directly explain the microcephaly phenotype: ‘mitotic cell cycle’ and ‘DNA replication’. We therefore considered these as core modules of the network. It is of note that the remaining 7 modules covered a broad range of biological processes and showed a varying degree of connection to the core modules, indicating functional relationship (Figures 3I and S4D). Interestingly the proportion of direct PRC2 target genes in each module was also diverse (Figure S4 E). The core modules contained the least proportion of direct PRC2 targets, whereas synapse and Ca^2+^ signaling modules contained a high number of direct PRC2 target genes (Figures 3I, Figure S4 E). In summary we concluded that a subset of highly connected cKO-*Eed*-MADM specific DEGs were sufficient to explain the microcephaly phenotype.

### Whole tissue but not sparse KO of *Eed* leads to progenitor proliferation defects through deregulation of multiple redundant gene modules

The STRING network suggested DNA replication/cell cycle as core modules that were deregulated due to tissue-wide PRC2 deficiency. To directly test this possibility we investigated progenitor proliferation potential in control-MADM, *Eed*-MADM and cKO-*Eed*-MADM. We performed 24h EdU pulse-chase experiments by injecting EdU into pregnant dams at E14.5 and analyzed the developing cortex at E15.5. Quantification of EdU^+^/GFP^+^ MADM-labelled cells showed that green RGPs in *Eed*-MADM (*Eed^-/-^*) displayed similar proliferation rates like green RGPs in control-MADM (*Eed^+/+^*) (Figures 4A-F, 4J). In contrast, green RGPs from cKO-*Eed*-MADM (*Eed^-/-^*) cortices showed 2.5fold reduction in EdU incorporation (Figures 4G-J). Thus, our results confirm the findings from the STRING network analysis and identify progenitor proliferation as a core module in the microcephaly phenotype.

**Figure 4.**
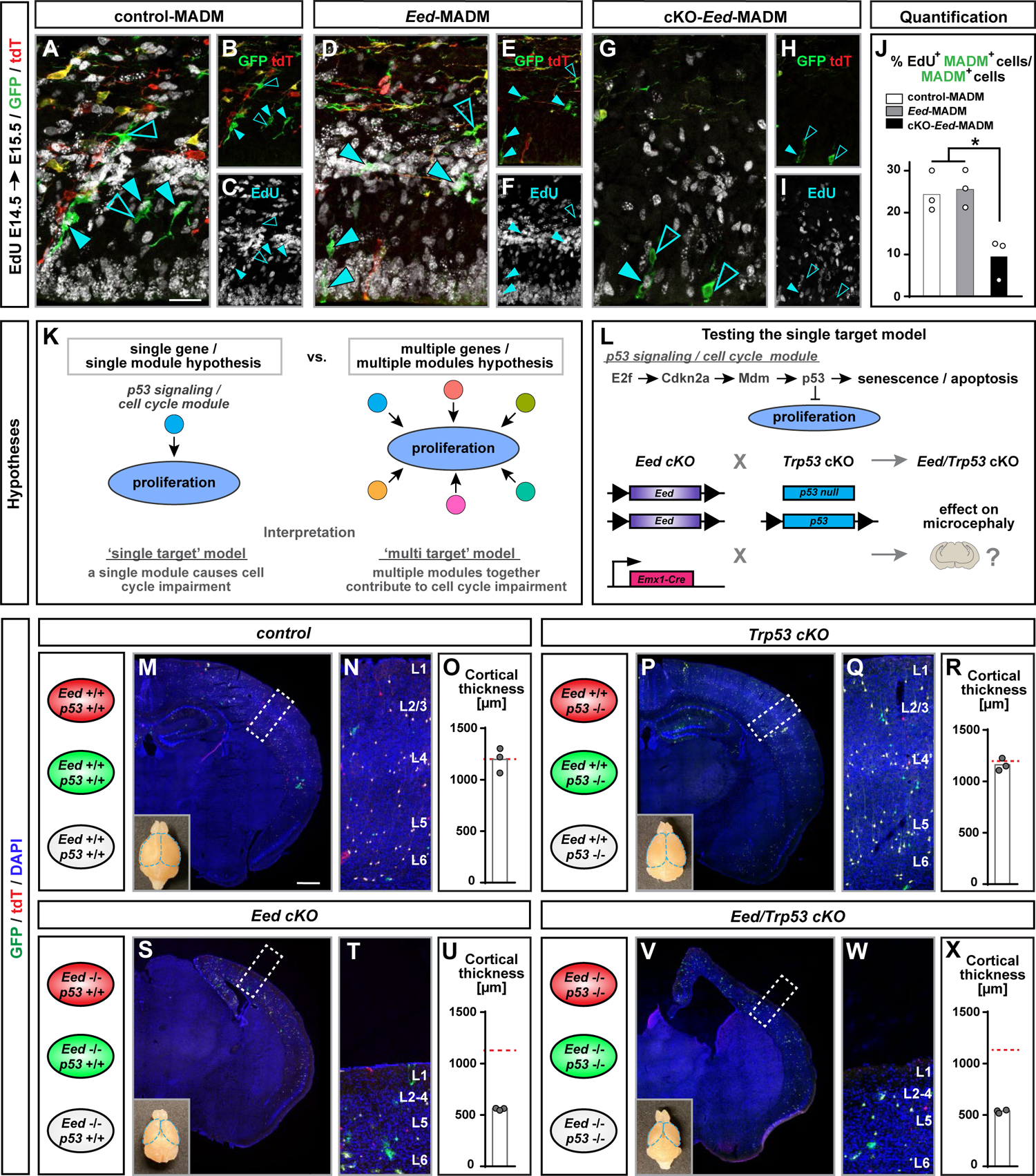
Proliferation deficits in *Eed^-/-^* RGPs upon global tissue-wide loss of PRC2 activity occur independently of *Trp53* expression. See also Figure S5. **(A-I)** Immunofluorescence images of EdU staining after a 24h EdU pulse administered at E14.5 to (A-C) control-MADM; (D-F) *Eed*-MADM; and (G-I) cKO-*Eed*-MADM mice. **(J)** Percentage of EdU^+^ green MADM-labelled cells in (white) control-MADM; (grey) *Eed*-MADM; and (black) cKO-*Eed*-MADM. **(K)** Schematic summary of two hypotheses arising from STRING network analysis. **(L)** Schematic illustration of experimental approach to address the single module hypothesis by creating *Eed/Trp53* double cKO in order to genetically inactivate the p53 signaling / cell cycle module. **(M-X)** Schematics of cellular genotypes in respective experimental MADM paradigms; and analysis of cortical thickness in (M-O) control, (P-Q) *Trp53* cKO, (S-U) *Eed* cKO, and (V-X) *Eed/Trp53* double cKO mice at P21. (M, P, S, V) Overview of MADM-labeling pattern in somatosensory cortex. (N, Q, T, W) depict higher magnification images of boxed areas in (M), (P), (S) and (V) with indications of cortical layers. Lower left insets in (M), (P), (S) and (V) show macrographs of whole brains with respective genotypes at P21. (R, U, X) Quantification of cortical thickness in (O) control; (R) *Trp53* cKO; (U) *Eed* cKO; and (X) *Eed/Trp53* double cKO mice. Each individual data point in (J), (O), (R), (U) and (X) represents one experimental animal. Statistics: (J) one-way ANOVA with multiple comparisons; * p<0.05; ** p<0.01; *** p< 0.001. Data indicate mean ± SEM. Scale bars: 25µm in (A, D, G); and 50µm in (B, C, E, F, H, I). 500µm in (M, P, S, V) and 60µm in (N, Q, T, W).

The topology of STRING network suggested a complex relationship of multiple deregulated gene modules culminating in proliferation defects. Following from this topology it was possible that multiple modules operated redundantly (multiple module hypothesis, Figure 4K left). Such model implied that removal of a single module would not alter the overall phenotype. Alternatively it was possible that a single module had a dominant effect on the phenotype (single module hypothesis, Figure 4K right). Removal of such predominant module was thus expected to rescue the microcephaly phenotype. Multiple lines of evidence supported the single module hypothesis. For example, a number of genetic mutations in human and mouse lead to microcephaly due to cell cycle dysregulation and apoptosis. Defects in cell cycle can originate from the impairment of a plethora of different biological processes, including DNA replication that we also identified as a core module in the STRING network. Despite the diverse genetic causes, microcephaly phenotypes often converge to trigger p53 activation and apoptosis in dividing RGPs and/or nascent neurons. In such cases genetic deletion of *Trp53* rescue the microcephaly phenotype (Bianchi et al., 2017; Breuss et al., 2016; Houlihan and Feng, 2014; Little and Dwyer, 2019; Mao et al., 2016; Marjanovic et al., 2015; Phan et al., 2021). Interestingly we noted the presence of significantly enriched p53-related GO terms in several modules of the STRING network (Supplementary Table S1). We thus considered p53 signaling as a strong candidate for the single module hypothesis (Figure 4L).

To directly assay for apoptosis we performed active CASPASE-3 stainings at E14.5 to determine the level of cell death. We detected a significant increase in the number of apoptotic cells (independent of MADM labelling) in cKO-*Eed*-MADM but not in *Eed*-MADM when compared to control-MADM and *Eed*-MADM cortices (Figures S5A-S5G). We concluded that global tissue-wide KO of *Eed*, and thus loss of PRC2 activity, not only translated to reduced proliferative potential in RGPs, but also increased apoptosis. In contrast, sparse or clonal ablation of *Eed* did not lead to RGP proliferation and/or survival deficits.

We proceeded with testing the ‘single target hypothesis’ by pursuing genetic epistasis experiments to inactivate the p53 signaling pathway in cKO-*Eed*-MADM cortices (Figure 4L). We utilized cKO-*Eed*-MADM paradigm and *Trp53*-deficiency models – *Trp53*-flox (Marino et al., 2000) and *Trp53* null allele (Jacks et al., 1994) – to generate conditional whole tissue (using *Emx1*-Cre) *Eed/Trp53* double mutants (cKO-*Eed*-MADM;*Trp53*-cKO abbreviated *Eed*/*Trp53* cKO); Figure 4L). We first confirmed that *Trp53* deletion abolished apoptosis (i.e. presence of CASPASE-3^+^ cells) in developing cortex in *Eed*/*Trp53* cKO mice (Figures S5I-5K). If microcephaly emerged in a predominantly p53-dependent manner in whole tissue *Eed* cKO, we expected a partial or full rescue of microcephaly in *Eed*/*Trp53* cKO. Cortical thickness and severity of microcephaly in both *Eed* cKO and *Eed/Trp53* cKO mice showed however similar reduction (Figures 4S-X) when compared to control and *Trp53* cKO, respectively (Figures 5M-R). We stained for CUX1 and noticed that in both *Eed* cKO and *Eed/p53* double cKO mice CUX1^+^ upper layer neurons were similarly reduced (Figures 5L-5P). Taken together our results indicated that elimination of the single *Trp53* gene module did not rescue the proliferation defect observed in *Eed* cKO mice. The data therefore supports the multiple module hypothesis, where microcephaly originates from multiple, functionally redundant, deregulated gene modules, only one of which being *Trp53*-dependent.

**Figure 5.**
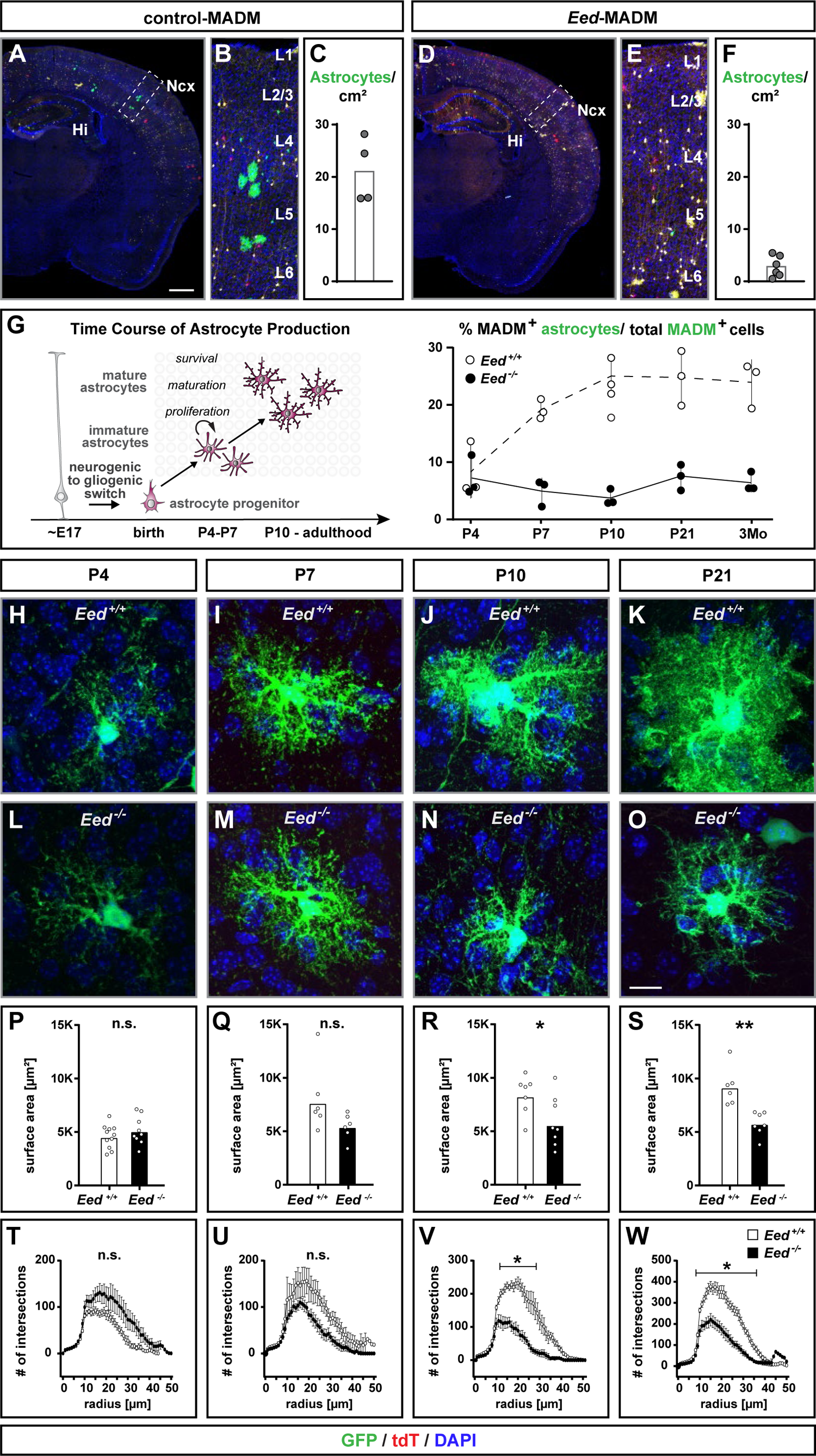
*Eed* is cell-autonomously required for cortical astrocyte production and maturation. See also Figure S6. **(A-F)** Overview of MADM-labeling pattern in the cortex in (A) control-MADM and (D) *Eed*-MADM mice at P21. (B, E) Boxed areas in (A) and (D) illustrating a higher resolution insets of the cortex with layer indications. (C, F) Quantification of astrocyte numbers with green MADM-labelled astrocytes/cm^2^ in (C) control-MADM; and (F) *Eed*-MADM mice. **(G)** Percentage of green astrocytes in control-MADM (white); and *Eed*-MADM (black) at P4, P7, P10, P21, and 3 months. **(H-O)** High resolution images of individual astrocytes from (H, I, J, K) control-MADM and (L, M, N, O) *Eed*-MADM. **(P-W)** Quantification of (P, Q, R, S) surface area; and (T, U, V, W) Sholl analysis of *Eed^+/+^* (white) and *Eed^-/-^* (black) astrocytes at (H, L, P, T) P4, (I, M, Q, U) P7, (J, N, R, V) P10, and (K, O, S, W) P21. (F, G): Each individual data point represents one experimental animal. Data show mean ± SEM. (P-W): Each data point represents one individual astrocyte. Data show mean ± SEM. Statistics for (P, Q, R, S) surface area: unpaired t-test with * p<0.05; ** p<0.01; *** p<0.001. Statistics for (T, U, V, W) Sholl analysis: multiple t-tests using the Two-stage linear step-up procedure of Benjamini, Krieger and Yekutieli with Q=1%; * p<0.05; ** p<0.01; ***p< 0.001. Scale bars: 500µm in (A and D); 60µm in (B and E); and 10µm in (H-O).

### PRC2 cell-autonomously regulates astrocyte production and maturation in developing neocortex

Upon neurogenesis RGPs obtain gliogenic potential and PRC2-mediated transcriptional repression has been implicated in cortical astrocyte generation, albeit controversial findings have been reported (Hirabayashi et al., 2009; Pereira et al., 2010). To address the role of PRC2 in gliogenic RGP lineage progression we utilized mosaic *Eed*-MADM enabling the assessment of the cell-autonomous *Eed* function at single cell level. We first determined whether PRC2 is active in developing astrocytes by performing H3K27me3 staining in immature astrocytes in control-MADM and *Eed*-MADM at P4. We found that developing astrocytes in control-MADM mice showed pronounced H3K27me3 staining while green *Eed^-/-^* astrocytes in *Eed*-MADM cortices showed no detectable H3K27me3 signal (Figures S6A-S6D).

In a first analysis we quantified absolute numbers (per cm^2^) of mature cortical MADM-labeled green (GFP^+^) astrocytes in control-MADM (*Eed^+/+^*) and *Eed*-MADM (*Eed^-/-^*) at P21. We observed significantly reduced numbers of *Eed^-/-^* astrocytes in *Eed*-MADM when compared to *Eed^+/+^* astrocytes in control-MADM (Figures 5A-5F). Next we performed time course analysis (Figure 5G, left) to determine whether reduced numbers of *Eed^-/-^* astrocytes in *Eed*-MADM arise from an impairment in (1) neurogenic-to-gliogenic switch in RGPs; (2) proliferation of astrocyte progenitors; or (3) survival of astrocytes. We quantified GFP^+^ astrocytes in control-MADM and *Eed*-MADM mice at ages P4, P7, P10, P21 and 3 months (Figure 5G, right). We detected comparable numbers of *Eed^+/+^* and *Eed^-/-^* nascent astrocytes at P4, indicating that the neurogenic-to-gliogenic switch in RGPs was not compromised upon loss of PRC2 activity. However, *Eed^+/+^* astrocytes underwent substantial expansion by proliferation in the first two postnatal weeks, indicated by increasing astrocyte numbers from P4 to P10. This observation was in agreement with recent findings (Clavreul et al., 2019), using lineage tracing at clonal level to show that astrocytes mostly expand during the first postnatal week. In contrast, the proportion of *Eed^-/-^* astrocytes remained rather constant from P4 onward and throughout the entire time course until 3 months (Figure 5G, right). Thus PRC2 cell-autonomously controls proliferation in astrocyte progenitors and/or promotes the survival rate of immature astrocytes.

Previous studies have shown that astrocytes undergo a morphological maturation phase from the first until the third postnatal week (Yang et al., 2013). We thus performed high-resolution imaging of individual green *Eed^+/+^* and *Eed^-/-^* astrocytes in time course analysis (Figures 5H-5O). We determined the total surface area (Figures 5P-5S); and extent of astrocyte process branching using Sholl analysis (Figures 5T-5W). While the surface area and ramification of processes in *Eed^+/+^* and *Eed^-/-^* astrocytes were comparable at P4 and P7 (Figures 5P-Q and 5T-U), we found that *Eed^-/-^* astrocytes showed significant impairment in surface expansion and ramification of complex processes from P10 onward (Figures 5R-S, 5V-W). We concluded that cell-autonomous PRC2 activity, besides controlling cortical astrocyte production, also fulfilled essential function in astrocyte maturation and branching.

### Transcriptomic signature of sparse immature *Eed^-/-^* astrocytes correlates with their reduced proliferation potential

In order to more comprehensively assess the phenotype of *Eed^-/-^* astrocytes, we utilized a strategy that allowed us to isolate ultrapure astrocyte populations in MADM context (Laukoter et al., 2020a; Laukoter et al., 2020c). In brief, we crossed *Eed*-MADM mice with transgenic *lacZ* reporter mice [*XGFAP-lacZ*; (Brenner et al., 1994)], expressing *lacZ* under the control of the human *GFAP* promoter and thus highly specifically in cortical astrocytes. By using FACS we purified MADM-labeled astrocytes (GFP^+^/lacZ^+^) and neurons (GFP^+^/lacZ^-^) at P4, when the numbers and morphology of green *Eed^+/+^* and *Eed^-/-^* astrocytes were comparable, and subjected respective samples to RNAseq (Figure 6A, Supplementary Table S2). We confirmed the purity of isolated neuron and astrocyte populations by probing for the expression of specific cell identity markers. As expected, *Aqp4* and *Gfap* were highly enriched in astrocyte populations while expression of *Lrrn3* and *Npm1* was much higher in neuron populations, independent of the *Eed* genotype (Figures S6E-S6F). Next we validated efficient *Eed* deletion in *Eed^-/-^* FACS-isolated astrocytes (Figure S6 G). In order to assess whether loss of PRC2 activity impacts astrocyte identity, we analyzed the expression of a previously defined set of astrocyte-specific genes (Bayraktar et al., 2020) in *Eed^+/+^* and *Eed^-/-^* astrocytes, respectively. We did not find consistent differences in marker gene expression for different astrocyte populations in *Eed^+/+^* and *Eed^-/-^* astrocytes. Thus *Eed*-deficiency seems to not alter astrocyte cell identity (Figure S6 H). Upon comparison of the transcriptional profiles in *Eed^+/+^* and *Eed^-/-^* astrocytes, we identified 317 DEGs, of which the majority was downregulated (Figure 6B). Gene set enrichment analysis (GSEA) identified mitotic cell cycle process as a significantly enriched GO term with a majority of genes being downregulated (Figures 6C-6D). These results imply a cell-autonomous proliferation phenotype in *Eed^-/-^* astrocytes.

**Figure 6.**
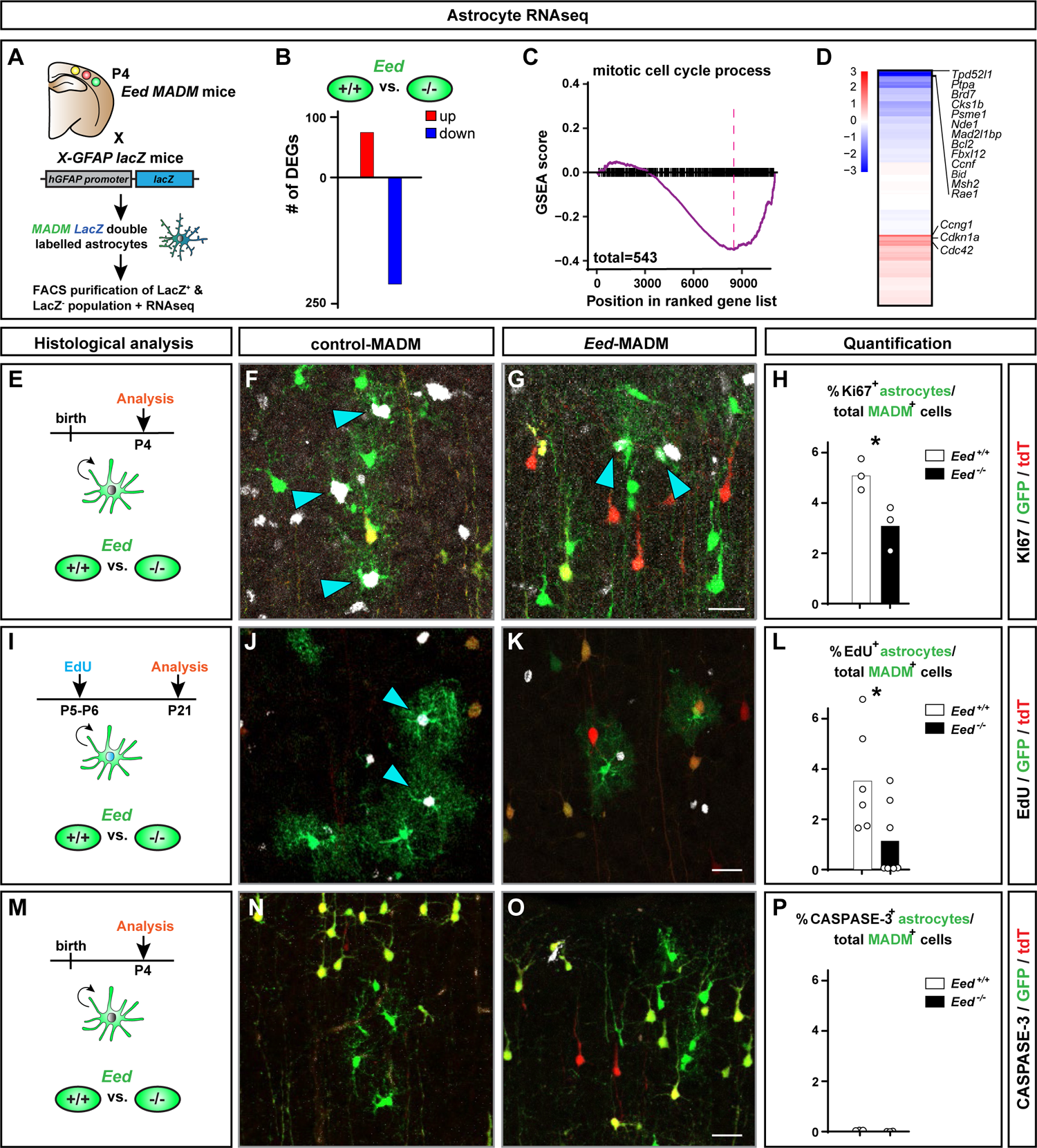
Cell-autonomous loss of *Eed* results in DEGs associated with mitotic cell cycle process, proliferation deficits but no increase in apoptosis in developing cortical astrocytes. See also Figure S6. **(A)** Schematic overview of genetic strategy using the *X-GFAP lacZ* transgene in combination with MADM to isolate highly pure GFP^+^/lacZ^+^ astrocytes by FACS at P4 for RNAseq. **(B)** Number of DEGs (padj < 0.2, DESeq2) upon comparison of green *Eed*^-/-^ (*Eed*-MADM) and *Eed*^+/+^ (control-MADM) astrocytes. **(C)** Running score plot for the term *mitotic cell cycle process* determined by Gene Set Enrichment Analysis (GSEA). **(D)** Heat map representing gene expression of cell cycle genes in the term *mitotic cell cycle process*. Selected genes are indicated. **(E-H)** Proliferation analysis of cortical astrocytes at P4 using KI67 staining. (E) Schematic overview of the experimental setup. (F-G) Representative images of GFP^+^ astrocytes in (F) control-MADM and (G) *Eed*-MADM. (H) Percentage of KI67^+^ proliferating *Eed*^+/+^ (white) and *Eed*^-/-^ (black) astrocytes. **(I-L)** Proliferation analysis of cortical astrocytes using EdU incorporation. (I) Schematic overview of the experimental setup with EdU injection at P5/P6 and analysis at P21. (J-K) Representative images of GFP^+^ astrocytes in (J) control-MADM and (K) *Eed*-MADM (L). Percentage of EdU^+^ proliferating *Eed*^+/+^ (white) and *Eed*^-/-^ (black) astrocytes. **(M-P)** Cell death analysis of cortical astrocytes using CASPASE-3 stainings. (M) Schematic overview of the experimental setup with CASPASE-3 stainings in P4 cortex. (N-O) Representative images of GFP^+^ astrocytes in (N) control-MADM and (O) *Eed*-MADM. (P) Percentage of *Eed*^+/+^ (white) and *Eed*^-/-^ (black) CASPASE-3^+^ astrocytes. Each individual data point represents one experimental animal. Statistics: unpaired two-tailed t-test with * p<0.05; ** p<0.01; *** p<0.001. Data indicate mean ± SEM. Scale bars: 20µm in (F and G); 50µm in (J and K); and 25µm in (N and O).

In order to determine the biological significance of our RNAseq analysis, we evaluated the levels of proliferation in *Eed^+/+^* and *Eed*^-/-^ astrocytes *in vivo*. First, we assessed the percentage of Ki67^+^ green *Eed^+/+^* and *Eed^-/-^* astrocytes at P4 (Figure 6E). We found a significant reduction in the proportion of Ki67^+^ *Eed^-/-^* astrocytes when compared to *Eed^+/+^* astrocytes (Figures 6F-6H). Next, we performed EdU injections into pups at age P5-P6 and analyzed the percentage of EdU^+^ green *Eed^+/+^* or *Eed^-/-^* astrocytes at P21 (Figure 6I). We found reduced amounts of EdU^+^ *Eed^-/-^* astrocytes when compared to *Eed^+/+^* astrocytes (Figures 6J-6L). To evaluate the level of apoptosis (potentially reducing numbers of *Eed^-/-^* astrocytes) we stained for activated CASPASE-3 at P4 (Figure 6M) but did not find any apoptotic figures in *Eed^+/+^* and *Eed^-/-^* astrocytes (Figures 6N-6P). Altogether, our results demonstrate cell-autonomous requirement of *Eed* and thus PRC2 activity in cortical astrocyte production.

## DISCUSSION

In this study we genetically dissected the cell-type-specific cell-autonomous and global tissue-wide PRC2 functions in RGP lineage progression during neocortical development with unprecedented single cell resolution. We utilized sparse and global *Eed* (essential component of PRC2) KO in combination with single cell MADM-labeling paradigms. Against our prediction based upon earlier work (Hirabayashi et al., 2009; Pereira et al., 2010; Telley et al., 2019) we found that PRC2 activity is not cell-autonomously required in cortical neurogenesis but exerts critical functions rather at the global tissue-wide level. In contrast, H3K27me3, catalyzed by PRC2, is cell-autonomously required for cortical astrocyte production and maturation. Altogether, our MADM-based analysis revealed distinct and sequential *Eed*/PRC2 functions in RGP lineage progression (Figure 7). Below, we discuss PRC2 requirement in cortical neurogenesis and astrocyte production in the context of individual cell-autonomous gene function and the interplay with tissue-wide genetic and cellular landscape.

**Figure 7.**
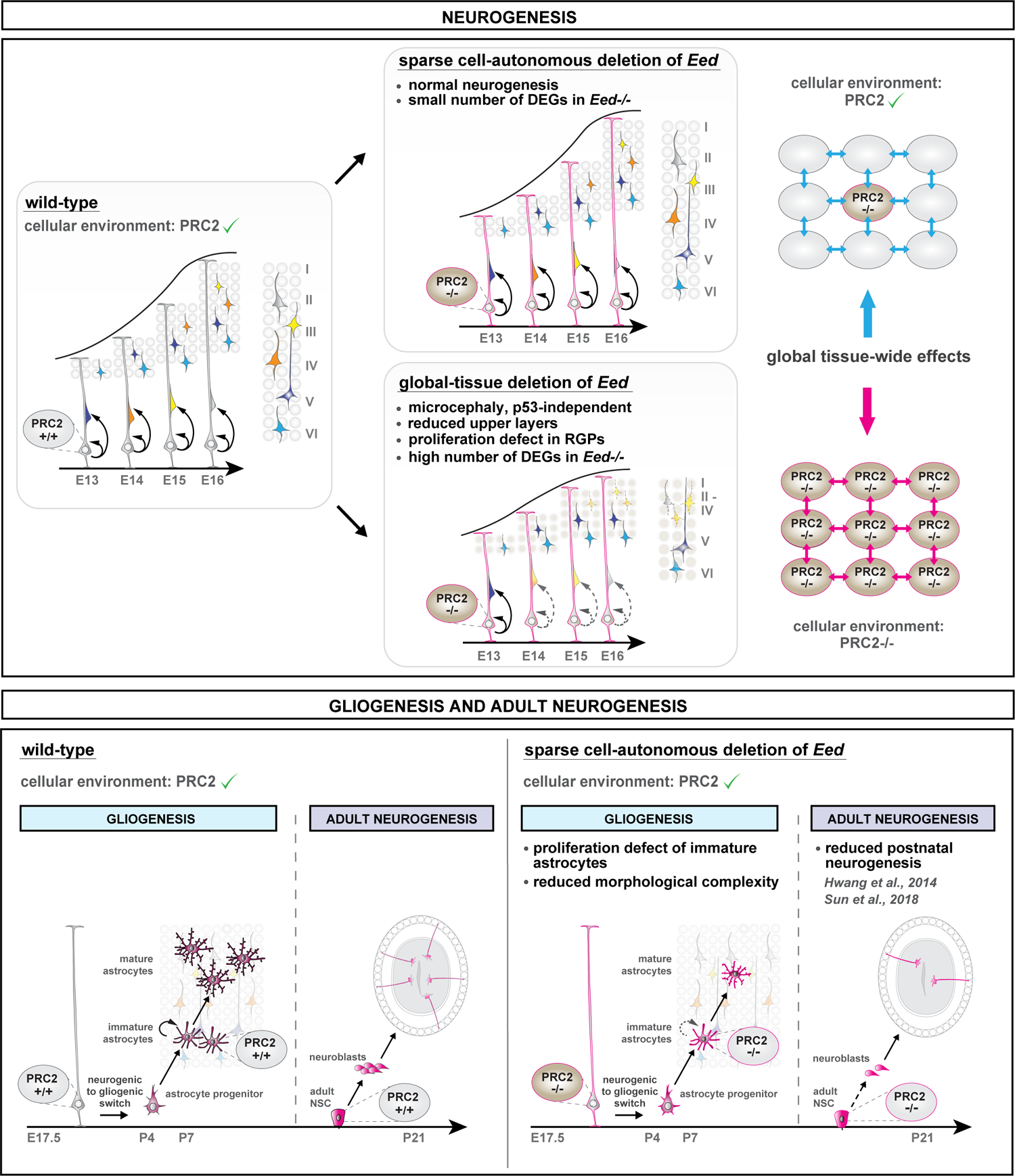
Distinct and sequential functions of PRC2 in neural stem cell lineage progression. Schematic model summarizing the findings of the study. PRC2 function is highly sensitive to the cellular environment during the neurogenic period. PRC2 is not cell-autonomously required to control cortical neurogenesis. Upon completion of RGP-mediated neurogenesis PRC2 cell-autonomously regulates astrocyte production and maturation. **(Top)** Individual *Eed^-/-^* RGPs in *Eed*-MADM show a small number of deregulated genes (DEGs) and no deficits in cortical neurogenesis. In contrast, RGPs in cKO-*Eed*-MADM display a high number of DEGs and proliferation defects (*Trp53*-independent) with diminished formation of upper layer neurons. Thus global tissue–wide loss of *Eed* triggers non-cell-autonomous effects and leads to microcephaly. **(Bottom)** *Eed*/PRC2 is cell-autonomously required in postnatal brain development for astrocyte production and maturation; and adult neurogenesis. In line with previous studies (Sun et al., 2018) and (Hwang et al., 2014), the sparse loss of *Eed* results in reduced postnatal neurogenesis with lower numbers of *Eed*^-/-^ granule cells in olfactory bulb.

### PRC2-dependent regulation of cortical neurogenesis and the relevance of global tissue-wide genetic and cellular environment

Using MADM, we genetically dissected the level of cell-autonomy of PRC2 function and the contribution of non-cell-autonomous tissue-wide genetic and cellular landscape in RGP lineage progression. We demonstrated that the sparse loss of *Eed* in an individual RGP surrounded by phenotypically normal cells did not affect embryonic neurogenesis. In contrast, when *Eed* gene function was ablated in an all mutant environment RGPs were compromised in their capacity to produce the correct number of cortical projection neurons. As a consequence, *Eed* cKO mice showed dramatic microcephaly. The distinct histological phenotypes of *Eed^-/-^* mutant cells in *Eed*-MADM (normal cortex size) and cKO-*Eed*-MADM (microcephaly) were mirrored at the global transcriptome level. The mutant cellular environment in cKO-*Eed*-MADM developing cortex induced a unique gene expression pattern in *Eed^-/-^* cells which was highly distinct from the one in *Eed^-/-^* cells upon sparse ablation.

Our data supports a two-step process upon loss of PRC2 activity leading to the terminal observed *Eed^-/-^* phenotype in cKO-*Eed*-MADM mice. First, PRC2-defiency triggers the upregulation of a relatively small set of direct target genes, which in the second step induce deregulation of a much larger number of indirect and/or secondary target genes. The concerted net result of direct and indirect deregulated gene expression in cKO-*Eed*-MADM ultimately results in impaired RGP proliferation dynamics, elevated apoptosis and global microcephaly. Two scenarios could explain the distinct phenotypic manifestation upon sparse and global KO of PRC2 activity. First, the heterozygous *Eed^+/-^* environment in *Eed*-MADM (sparse deletion) rescues RGPs from proliferation deficits by preventing exuberant deregulation of gene expression due to protective global-tissue wide effects. Second, the concerted loss of PRC2 activity in all cells in cKO-*Eed*-MADM triggers an exacerbated tissue-wide response that ultimately results in much increased ectopic expression of direct PRC2 target genes thereby inducing secondary waves of ectopic transcription leading to the specific deregulation of genes controlling cell cycle and apoptosis. Regardless of the precise mechanism it becomes clear that the genetic and cellular environment is critical for RGP lineage progression and the generation of correctly sized cortex (Figure 7, top).

### RGP proliferation deficits and microcephaly due to global PRC2 KO occurs independent of *Trp53* expression

Gene network analysis of DEGs specific for cKO-*Eed*-MADM has uncovered particular modules comprised of functionally related genes. The core modules encompass proliferation-associated genes. Many primary microcephaly mutants show a strong mechanical impairment of cell division (defective assembly of the mitotic spindle or centromere attachment) (Bianchi et al., 2017; Breuss et al., 2016; Keil et al., 2020; Little and Dwyer, 2019; Mao et al., 2016; Marino et al., 2000; Marjanovic et al., 2015; Nechiporuk et al., 2016; Phan et al., 2021) and a strong correlation between cell cycle mechanisms involved in DNA integrity, mitotic checkpoints and prevention of DNA damage-induced apoptosis via the p53 pathway. Recent studies have unraveled a similar function for the chromatin remodelers Ino80 and REST. Both remodelers were shown to prevent accumulation of DNA damage and associated p53-dependent microcephaly in the forebrain (Keil et al., 2020; Nechiporuk et al., 2016).

Based on genetic epistasis experiments we demonstrated that diminished RGP proliferation and associated microcephaly in *Eed* cKO mice could not be rescued by concomitant loss of *Trp53*. These findings point towards a *Trp53*-independent mechanism and are insofar intriguing since other gene mutations affecting cell cycle progression trigger microcephaly in a *Trp53*-dependent manner (Bianchi et al., 2017; Breuss et al., 2016; Keil et al., 2020; Little and Dwyer, 2019; Mao et al., 2016; Marino et al., 2000; Marjanovic et al., 2015; Nechiporuk et al., 2016; Phan et al., 2021). Given that the global mutant cellular landscape in cKO-*Eed*-MADM mice is the main driver of microcephaly, it is likely that the brain size alteration due to whole-tissue loss of PRC2 activity involves a complex aggregation or interplay of environmental cues and cell-intrinsic transcriptional responses. The systemic environment-dependent reciprocity of cellular behavior in global *Eed* mutant cortex may be similar to the concept of a synthetic defect (i.e. a fitness defect upon whole-tissue loss of PRC2 activity in all cortical progenitor cells). Interestingly, a recent study employing transformed mouse embryonic fibroblasts (MEFs) revealed a PRC2-dependent mechanism to control cell proliferation independent of *Cdkn2a*-*p53*-*pRB* (Piunti et al., 2014). The authors provide evidence that Polycomb complexes actually localize at replication forks to direct progression and symmetry of DNA replication, thereby controlling cell proliferation independent of major cell cycle checkpoint genes and the p53 pathway. It is however not clear, how cell cycle progression by Polycomb localization to replication forks is controlled at the individual cell level and which role the genetic landscape of the cellular environment plays in this process.

### Cell-autonomous function of PRC2 in astrocyte development

A complex interplay of repressive mechanisms (e.g. DNA- and histone methylation) and activating mechanisms (e.g. histone acetylation and chromatin remodeling) regulates gliogenesis and the fate choice between astrocyte and oligodendrocyte identity (Fan et al., 2005; Hatada et al., 2008; Hirabayashi et al., 2009; Lattke et al., 2021; Lessard et al., 2007; Namihira et al., 2009; Pereira et al., 2010; Sun et al., 2018; Takizawa et al., 2001; Tan et al., 2012; Wang et al., 2020).

Based on global-tissue deletion studies, PRC2 was previously thought to play an essential role in regulating the neurogenic-to-gliogenic switch of RGPs (Hirabayashi et al., 2009; Pereira et al., 2010). However, sparse cell-autonomous *Eed* deletion revealed that lineage progression of neurogenic RGPs to astrocyte intermediate progenitor cells (aIPCs) was unaffected, both in terms of numbers as well as molecular markers. These results underline that astrocyte specification following the neurogenic-to-gliogenic switch is not impaired by PRC2-deficiency at the single cell level.

Following aIPC identity acquisition, cortical astrocytes are reported to scatter throughout all cortical layers while exhibiting a high proliferation rate during the first postnatal week (P0-P7) (Clavreul et al., 2019). This dynamic dispersion and proliferation phase gradually progresses into a maturation phase (P7–P21) where individual astrocytes increase in volume and process complexity (Clavreul et al., 2019). While many studies have addressed the function of epigenetic regulation during the onset of gliogenesis (Fan et al., 2005; Hatada et al., 2008; Hirabayashi et al., 2009; Lattke et al., 2021; Lessard et al., 2007; Namihira et al., 2009; Pereira et al., 2010; Sun et al., 2018; Takizawa et al., 2001; Tan et al., 2012; Wang et al., 2020), the role of epigenetics in controlling proliferation and maturation of immature astrocytes has not been analyzed in detail yet. Based on the single cell resolution provided by our MADM approach, we were able to assess the consecutive steps of astrocyte development both quantitatively and qualitatively. We thereby uncovered an essential function of PRC2 in aIPCs: despite *Eed^-/-^* aIPCs were correctly specified PRC2-deficiency resulted in profound proliferation and maturation defects (Figure 7, bottom).

### The implication of distinct and sequential PRC2 function in RGP lineage progression

Our findings add to several lines of evidence implying that sparse alteration of gene function at the individual cell level affects astrocyte production, but is not cell-autonomously required for neurogenesis (Beattie et al., 2017; Laukoter et al., 2020c). However, certain genes only cause a significant disruption of faithful neurogenesis upon deletion at the global tissue level (Beattie et al., 2017). The predominance of tissue-wide effects causing severe cortical malformations has recently also been reported for genes involved in migration or cell polarity (Hansen et al., 2022), thus further underlining that global tissue or non-cell-autonomous systemic effects significantly shape the early steps of cortical development such as neuron production (Hansen and Hippenmeyer, 2020).

The discovery of distinct and sequential functions of PRC2 in lineage progression will have profound implications for our understanding of diseases arising from postzygotic mutations in individual dividing cells (i.e. somatic cell mosaicism). Somatic cell mosaicism can occur at any point throughout development and is associated with a number of disorders, including neurological diseases and tumorigenesis (D’Gama and Walsh, 2018; Frank, 2014; Jamuar et al., 2014). In the future, it will be of great relevance to investigate the contribution of the cellular environment (i.e. the niche) to cell-autonomous or non-cell-autonomous phenotype manifestations upon acquisition of mutations altering the function of particular epigenetic mechanisms. Somatic disease-promoting mutations altering the repressive mechanisms mediated by PRC2 may either affect the expression of PRC2 core components (such as *Ezh2* overexpression (Suvà et al., 2009) or PRC2 recruitment to chromatin [such as H3K27M mutations (Bender et al., 2013; Harutyunyan et al., 2019; Lewis et al., 2013)]. Such mutations will be interesting candidates for the assessment of disease development originating from a single PRC2 mutant stem cell in a cell type- and environment-dependent manner.

## ACKNOWLEDGEMENTS

We thank A. Heger (IST Austria Preclinical Facility), A. Sommer and C. Czepe (VBCF GmbH, NGS Unit) and S. Gharagozlou for technical support. This research was supported by the Scientific Service Units (SSU) of IST Austria through resources provided by the Imaging & Optics Facility (IOF), Lab Support Facility (LSF), and Preclinical Facility (PCF). N.A. received funding from the FWF Firnberg-Programm (T 1031). The work was supported by IST institutional funds and by the European Research Council (ERC) under the European Union’s Horizon 2020 research and innovation program (grant agreement 725780 LinPro) to S.H.

## Author Contributions

N.A. and S.H. conceived the project. N.A. performed wet lab experiments. F.M.P. performed bioinformatics analysis. C.S. provided help with mouse genetics and breeding. N.A., F.M.P. and S.H. wrote the original draft. All authors edited and revised the manuscript.

## MATERIAL AND METHODS

### Mouse Lines

All mouse colonies were maintained in accordance with protocols approved by institutional animal care and use committee, institutional ethics committee and the preclinical core facility (PCF) at IST Austria. Experiments were performed under a license approved by the Austrian Federal Ministry of Science and Research in accordance with the Austrian and EU animal law (license number: BMWF-66.018/0007-II/3b/2012 and BMWFW-66.018/0006-WF/V/3b/2017). Mice with specific pathogen free status according to FELASA recommendations (Mahler Convenor et al., 2014) were bred and maintained in experimental rodent facilities (room temperature 21 ± 1°C [mean ± SEM]; relative humidity 40%-55%; photoperiod 12L:12D). Food (V1126, Ssniff Spezialitäten GmbH, Soest, Germany) and tap water were available *ad libitum*.

Mouse lines with MADM cassettes inserted in Chr. 7 (Hippenmeyer et al., 2013), *Eed*-flox (Yu et al., 2009), *Trp53*-flox (Marino et al., 2000), *Trp53* null (Jacks et al., 1994), *hGFAP-lacZ* (Brenner et al., 1994), *Emx1*-Cre (Gorski et al., 2002), and *Emx1*-Cre^ERT2^ (Kessaris et al., 2006), have been described previously.

Calculation of recombination probability and recombination of *Eed-flox* allele onto chromosomes carrying the MADM cassettes was performed according to standard techniques. These are described in detail elsewhere (Amberg and Hippenmeyer, 2021) and are summarized in Figure S1. All MADM-based analyses were carried out in a mixed C57BL/6, CD1 genetic background, in male and female mice without sorting experimental cohorts according to sex. No sex specific differences were observed under any experimental conditions or in any genotype. Based on genotype, experimental groups were randomly assigned. The age of experimental animals is indicated in the respective figures, figure legends and source data files.

### Tissue Isolation and Immunostaining

From P4 onwards, mice were deeply anesthetized by injection of a ketamine/ xylazine/ acepromazine solution (65 mg, 13 mg and 2 mg/kg body weight) and unresponsiveness was confirmed through pinching the paw. The diaphragm of the mouse was opened from the abdominal side to expose the heart. Cardiac perfusion was performed with ice-cold PBS (phosphate-buffered saline) followed immediately by 4% PFA (paraformaldehyde) prepared in PB buffer. Brains were removed and further fixed in 4% PFA O/N to ensure complete fixation. Brains were cryopreserved with 30% sucrose (Sigma-Aldrich) solution in PBS for approximately 48 h and were then embedded in Tissue-Tek O.C.T. (Sakura).

Pregnant females were sacrificed at the respective time points to obtain E12, E13, E14, E15 and E16 embryonic brain tissue. Embryonic and P0 brains were directly transferred into ice-cold 4% PFA. Cryopreservation was performed in 30% sucrose in PBS and OCT embedding was performed according to standard techniques. For embryonic timepoints and postnatal P0, 20-25µm coronal frozen sections were directly mounted onto superfrost glass-slides (Thermo Fisher Scientific) and stored at −20°C until further usage.

For adult time points, 30-45 µm coronal frozen sections were collected in 24 multi-well dishes (Greiner Bio-one) and stored at −20 °C in antifreeze solution (30% v/v ethyleneglycol, 30% v/v glycerol, 10% v/v 0.244 M PO_4_ buffer) until used. Embryonic sections were removed from −20°C and allowed to acquire room temperature for 15min, followed by three wash steps (5 min) with PBS. Tissue sections were blocked for 30 min in a blocking buffer (containing 5% horse serum (Thermo Fisher Scientific) and 1% Triton X-100 in PBS). Primary antibodies were diluted in staining buffer (5% horse serum, 0.5% Triton X-100) and incubated overnight at 4 °C. Sections were washed three times for 5 min with PBS and incubated with corresponding secondary antibody diluted in staining buffer for 2h at RT. Sections were washed three times with PBS. Adult brain sections were transferred to a 24well-plate filled with PBS and stained as floating sections following the above mentioned steps. After the washes following the secondary antibody staining, sections were mounted onto superfrost glass-slides (Thermo Fisher Scientific) and allowed to dry. Nuclear staining of glass-mounted brain sections was performed by 10 min incubation with PBS containing 2.5% DAPI (4’,6-diamidino-2-phenylindole, Thermo Fisher Scientific). Sections were embedded in mounting medium containing 1,4-diazabicyclooctane (DABCO; Roth) and Mowiol 4-88 (Roth) and stored at 4 °C until imaging.

Primary antibodies: chicken anti-GFP (AVES, dilution 1:400), goat anti-mCherry (SICgen, dilution 1:400), rabbit anti-CTIP (Abcam, dilution 1:400), goat anti-CUX1 (Santa Cruz, dilution 1:100), rabbit anti-CASPASE-3 (Cell Signaling, dilution 1:500), rabbit anti-H3K27me3 (Diagenode, dilution 1:1,000).

Secondary antibodies: donkey anti-chicken-FITC secondary antibody (Invitrogen, dilution 1:500), donkey anti-goat Alexa568 secondary antibody (Molecular Probes, dilution 1:1,000), donkey anti-goat Alexa647 secondary antibody (Molecular Probes, dilution 1:1,000), donkey anti-rabbit Alexa647 secondary antibody (Molecular Probes, dilution 1:1,000).

### MADM Clonal Analysis

MADM clone analysis was performed as previously described (Beattie et al., 2020). In brief, timed pregnant females were injected intraperitoneally with tamoxifen (TM) (Sigma) dissolved in corn oil (Sigma) at E11 or E12 at a dose of 2-3 mg/pregnant dam. Live embryos were recovered at E18–E19 through caesarean section, fostered, and raised for further analysis at P21. For embryonic time point analysis, caesarean section and analysis was performed at either E13 or E16. Brains containing MADM clones were isolated, and tissue sections processed as described above.

### EdU Labeling Experiments

Cell-cycle experiments were based on the use of the Click-iT Alexa Fluor 647 imaging kit (Thermo Fisher C10340). Reagents were reconstituted according to the user manual. Pregnant females were injected with EdU (1mg/ml EdU stock solution; 50µl/10g mouse) at E14.5. Embryos were collected 24 hours after EdU injection. Tissue was fixed in 4% PFA and immunohistochemistry was performed as described above, except that the Click-iT imaging kit was used (according to the instruction manual) to visualize the EdU signal before performing the DAPI staining. P5/P6 pups were injected with EdU (1 mg/ml EdU stock solution; 30 µl/mouse). Mice were perfused at P21 using 4% PFA. Immunohistochemistry was performed as described above, except that the Click-iT imaging kit was used (according to the instruction manual) to visualize the EdU signal before performing the DAPI staining.

### Imaging and Quantification

Sections were imaged using an inverted LSM800 confocal microscope (Zeiss) and processed using Zeiss Zen Blue software. Confocal images were analyzed in Photoshop software (Adobe) by manually counting MADM-labeled cells based on respective marker expression (Figures 1-3 and Figures 5-6). To determine cortical thickness solely DAPI images were used. Images were opened in Zen Blue software and measurements were performed using the “line”-tool of this software. Three different measurements in the somatosensory cortex were done per image and accordingly combined to one value by averaging the three measured values (Figures 1 and 5). For cell quantification and cortical thickness measurements a minimum of six sections were analyzed per individual brain. Three different individuals were analyzed per genotype. Significance was determined using ANOVA test in Graphpad Prism 7.0. For clonal analysis (Figure 2), 2D clone reconstruction was performed by using a custom script in Image J (Beattie et al., 2020). Cortical areas were identified by using the Allen Brain Atlas (http://mouse.brain-map.org/static/atlas). Significance was determined using t-test in Graphpad Prism 7.0. For a detailed protocol please refer to (Beattie et al., 2020).

### Preparation of Single Cell Suspension and FACS

For embryonic tissue, pregnant females were sacrificed and E12/E13/E16 embryos were collected. For postnatal tissue, mice were sacrificed by decapitation (P0) or cervical dislocation (P4 onwards). Neocortex area was dissected. For MADM cortical bulk sorting, pools of 2-3 individual cortices were used to generate one sample. For astrocyte sorting, individual animals were used to generate replicates. Single-cell suspensions were prepared by using Papain containing l-cysteine and EDTA (vial 2, Worthington), DNase I (vial 3, Worthington), Ovomucoid protease inhibitor (vial 4, Worthington), EBSS (Thermo Fisher Scientific), DMEM/F12 (Thermo Fisher Scientific), FBS and HS. All vials from Worthington kit were reconstituted according to the manufacturer’s instructions using EBSS. The dissected brain area was directly placed into Papain-DNase solution (20 units/ml papain and 1000 units DNase). Enzymatic digestion was performed for 30 min at 37 °C in a shaking water bath. Next, solution 2 (EBSS containing 0.67 mg Ovomucoid protease inhibitor and 166.7 U/ml DNase I) was added, the whole suspension was thoroughly mixed and centrifuged for 5 min at 1000 rpm at RT. Supernatant was removed and cell pellet was resuspended in solution 2. Trituration with p1000 pipette was performed to mechanically dissolve any remaining tissue parts. DMEM/F12 was added to the cell suspension as a washing solution, followed by a centrifugation step of 5 min with 1500 rpm at RT. For MADM bulk sorting, cells were resuspended in media (DMEM/F12 containing 10% FBS and 10% HS) and kept on ice until sorted.

For astrocyte sorting, cells were incubated for 25min at 37°C in 100µl LacZ staining reagent (Abcam, diluted 1:50). Reaction was stopped by adding 100µl DMEM/F12 containing 10% FBS and 10% HS. Samples were centrifuged 5 min with 1500 rpm at RT and resuspended in DMEM/F12 containing 10% FBS and 10% HS. Samples were kept in the dark at room temperature until sorting was started. To ensure specificity of LacZ staining, a negative sample was always processed and sorting gates were adjusted accordingly. See further details at (Laukoter et al., 2020a).

Right before sorting, cell suspension was filtered using a 40 µm cell strainer. FACS was performed on a BD FACS Aria III using 100 nozzle and keeping sample and collection devices (0.8 ml PCR tubes) at 4 °C. Duplet exclusion was performed to ensure sorting of true single cells. Astrocytes and E12.5 MADM bulk cells were sorted into 4 µl lysis buffer (0.2% Triton X-100, 2U/µl RNase Inhibitor (Clonetech)), while E13.5, E16.5 and P0 MADM bulk samples were sorted into 50µl RNA extraction buffer 10mM EDTA pH 8.0, 30mM EDTA pH 8.0, 1% SDS, 200µm/µl Proteinase K in nuclease-free water). Immediately after sorting into 4µl lysis buffer was completed, samples were transferred into a 96-well plate (Bio-Rad) that was kept on dry ice. Once the plate was full it was sealed with AluminumSeal and kept at −80 °C until further processing.

### RNA Extraction and cDNA Library Preparation of MADM Samples for RNA Sequencing

After cell sorting of E13/E16 and P0 cells, samples were incubated for 30 min at 37 °C and stored at −80°C until further usage. After thawing samples on ice, their total volume was adjusted to 250 µl using RNase-free H_2_O (Thermo Fisher Scientific) followed by addition of 750 µl Trizol LS (Thermo Fisher Scientific) and inverting five times. After incubating for 5-min at RT, the entire solution was transferred into a MaXtract tube (Qiagen). 200µl microliters chloroform (Sigma-Aldrich) were added. After three times 5sec vortexing and 2min incubation at RT, samples were centrifuged for 2 min with 12,000 rpm at 18 °C. Supernatant was transferred to a new tube and isopropanol (Sigma-Aldrich) was added in a 1:1 ratio. For better visibility of the RNA pellet, 1 µl GlycoBlue (Thermo Fisher Scientific) was added and entire solution was mixed by vortexing 3 × 5sec. Samples were incubated at −20°C for 3 nights to precipitate RNA. Then, samples were centrifuged for 20min with 14,000 rpm at 4 °C. Supernatant was removed and RNA pellet was washed with 70% ethanol, followed by a 5min centrifugation step (14,000 rpm at 4 °C). RNA pellet was resuspended in 12.5 µl RNase-free H_2_O. RNA quality was analyzed using Bioanalyzer RNA 6000 Pico kit (Agilent) following the manufacturer’s instructions. RNA samples were pipetted into a 96well plate, sealed with AluminumSeal and stored at −80 °C until further use. Sequencing libraries were prepared following the Smart-Seq v2 protocol (Picelli et al., 2013) using custom reagents (VBCF GmbH) and libraries from a 96-well plate were pooled, diluted and sequenced on a HiSeq 2500 (Illumina) using v4 chemistry or NextSeq550 (Illumina).

## QUANTIFICATION AND STATISTICAL ANALYSIS

### Analysis of MADM-Labeled Brains

Sections were imaged using an inverted LSM800 confocal microscope (Zeiss) and processed using Zeiss Zen Blue software. Confocal images were imported into Photoshop software (Adobe) and MADM-labeled cells were manually counted based on morphology or respective marker expression as described previously (Beattie and Hippenmeyer, 2017; Beattie et al., 2017). Statistical analysis and data visualization was performed in Graphpad Prism 7.0.

### Astrocyte Morphology Analysis

Detailed analysis of astrocyte morphology was done as described previously (Beattie et al., 2017) with some modifications. Lower layer astrocytes that expressed GFP^+^ were imaged with a 63x oil objective and overlapping z-sections. Sholl analysis was performed to measure astrocyte branching complexity. 3D reconstruction and analysis was done using the Filament tracer algorithm of the IMARIS software. Total cell volume of astrocytes was assessed from the 3D structure.

### Processing and Analysis of Bulk RNA-seq Data from Embryonic Time Course

Read processing, alignment and annotations were described previously (Laukoter et al., 2020b). STAR (Dobin et al., 2013) alignment parameters: --outFilterMultimapNmax 1, -- outSAMstrandField intronMotif, --outFilterIntronMotifs RemoveNoncanonical and quantMode GeneCounts. Downstream analyses were performed in R (v3.6.1) unless indicated otherwise. Read counts of the deleted region in the *Eed* gene (chr7:89969506-89972357, mm10) were calculated using bedtools (Quinlan and Hall, 2010) intersect with the -split option on the aligned bam file produced by STAR.

We analyzed 118 samples and removed 29 samples with a low percentage of uniquely aligned reads (<60%), low absolute number of uniquely aligned reads (<1M) or due to low correlation with biological replicates. For E12.5 we also removed 2 samples due to high level of read counts in *Eed* deleted region indicating inefficient recombination.

**Figure S3B:** Read counts in *Eed* deleted regions were calculated as Reads Per Million total reads and plotted.

**Figure S3C:** For PCA plot varianceStabilizingTransformation with blind=T parameter was performed on raw read counts. Top 500 variable genes were identified using rowVars (matrixStats v0.56.0) and principal component analysis (PCA) was performed using prcomp.

**Figure 3E:** Statistics on differential expression between all pairs of genotypes were calculated with DESeq2 (v1.26) (Love et al., 2014) using contrasts for each developmental time point separately. To reduce noise only genes with an average read coverage of >1 (E12.5) or >10 (E13.5, E16.5, P0) were used in the analyses. We used an adjusted p-value (padj) cutoff of 0.1 for differentially expressed genes (DEG) and a log2FoldChange >0 (up-regulation) or <0 (down-regulation) for analyses.

**Figure S3D:** Score heatmap: All DEGs from all developmental time points were analyzed. For each gene the log10 of the uncorrected p-value, from the indicated comparison, was calculated and corrected to be positive for log2FoldChange > 0 and negative for log2FoldChange < 0. This score was cut at +5/-5 for better visualization. Heatmap was drawn with pheatmap package (v1.0.12).

**Figure 3G, S4B:** To determine the coverage of genes with H3K27me3 we used published data from (Albert et al., 2017). We downloaded Supplemental Table 2 (embj201796764-sup-0002-datasetev1.xlsx) and used columns 9-11 (aRG-P, aRG-N, bRG) for further analyses. Genes with duplicated Symbol IDs were removed. Direct target genes were defines as up regulated in the respective DEG analysis (log2FoldChange > 0) and reported to be marked in one or more of the above mentioned cell types by H3K27me3. Indirect target genes were defined as either down regulated or not marked by H3K27me3. Note that for this analysis we used DEGs (padj < 0.1) from *Eed*-MADM/control-MADM and cKO-*Eed*-MADM/control-MADM analyses that were also informative in (Albert et al., 2017).

**Figures 3I, S4C-D:** This analysis was performed with R v4.0.1. To identify protein protein interactions we used the STRING databse via the R package STRINGdb (v2.0.1). A STRINGdb object was initialised with STRINGdb$new and parameters: version=“11”, species=10090, score_threshold=600. We mapped the gene symbols of the 644 cKO-Eed-MADM specific DEGs to STRING IDs using the function map. 95% of genes could be mapped in this way. To identify the significance of the retrieved protein interaction network we used the function plot_network. The resulting network contained: 612 proteins with 1183 interactions. Expected interactions in at random: 681 and PPI enrichment p-value <10^-16^. Next we identified protein-protein interactions using the get_interactions function (STRINGdb). We created an igraph (v1.2.5) object using graph.data.frame with directed=F parameter, removed multiple edges and identified connected subnetworks using the function clusters. We extracted the largest connected network (333 genes) and repeated the above steps to create an igraph object with only these 333 genes. To visualize the network we exported the igraph object to Cytoscape (v3.7.2) (Gustavsen et al., 2019) using createNetworkFromIgraph function from RCy3 package (v2.8.0) (Shannon et al., 2003). We identified clusters of connected genes using get_clusters function (STRINGdb) with standard parameters. This analysis grouped the 333 genes in 9 clusters. Genes in each cluster were analysed using enrichGO (clusterProfiler package v3.16.0) with OrgDb = org.Mm.eg.db (v3.11.4), ont = “all”, readable = T, pool = T parameters. The top 3 enriched GO terms (Supplemental Table S1) were used to plot Figure S4C. Figure S4D: Edge thickness correlates to number protein-protein interactions between clusters.

### Processing and Analysis of Bulk RNA-seq Data from Purified Astrocytes

We analyzed 16 samples and removed 5 samples based on position on PCA plot or low correlation between biological replicates. All normalized gene expression values were identified using DESeq2’s counts function with normalize = T parameter. Heatmaps were drawn with pheatmap.

**Figure S6E and S6F:** For astrocyte marker genes the median normalized gene expression for neuron samples was set to 1 and the expression of astrocyte samples was plotted relative to that value. For neuron marker genes the median normalized expression for astrocyte samples was set to 1 and the expression of neuron samples was plotted relative to that value.

**Figure S6G:** *Eed* deletion counts were identified as described above and normalized expression values are shown.

**Figure S6H:** Normalized expression values of astrocyte markers identified by (Bayraktar et al., 2020) were averaged over all biological replicates. The highest expressing sample was set to 1 and expression of the respective other sample is shown relative to that sample.

**Figure 6B:** Statistics on differential expression between genotypes were calculated with DESeq2 (v1.26) (Love et al., 2014). We used an adjusted p-value (padj) cutoff of 0.2 for differentially expressed genes (DEG) and a log2FoldChange >0 (up-regulation) or <0 (down-regulation) for analyses.

**Figure 6C:** For gene set enrichment analysis a score was calculated as the log10 uncorrected p-value and corrected to be positive for log2FoldChange > 0 and negative for log2FoldChange < 0. DEGs were sorted based on this score and analysed using gseGO from the clusterProfiler package using ont = ‘BP’, OrgDb = org.Mm.eg.db, pvalueCutoff = 0.9, minGSSize = 500, maxGSSize = 1000 parameters. The term mitotic cell cycle process was the top enriched term with p-value: 0.011. Running scores were plotted using gseaplot from the clusterProfiler package.

**Figure 6D:** Score heatmap as described above with genes from GO term “mitotic cell cycle progress”, values cut at +3/-3.

## RESOURCE AVAILABILITY

### Lead Contact

Further information and requests for resources and reagents should be directed to and will be fulfilled by the Lead Contact, Simon Hippenmeyer.

### Materials Availability

All published and inaugural reported reagents and mouse lines will be shared upon request within the limits of the respective material transfer agreements.

### Data and Code Availability

Raw sequencing data and sample lists are accessible as super series at GEO under accession number GSE191109. All data have been presented in Figures and Supplemental Figures. Original images will be made available upon request.

## SUPPLEMENTARY MATERIAL

### Supplementary Figure Legends

**Figure S1.**
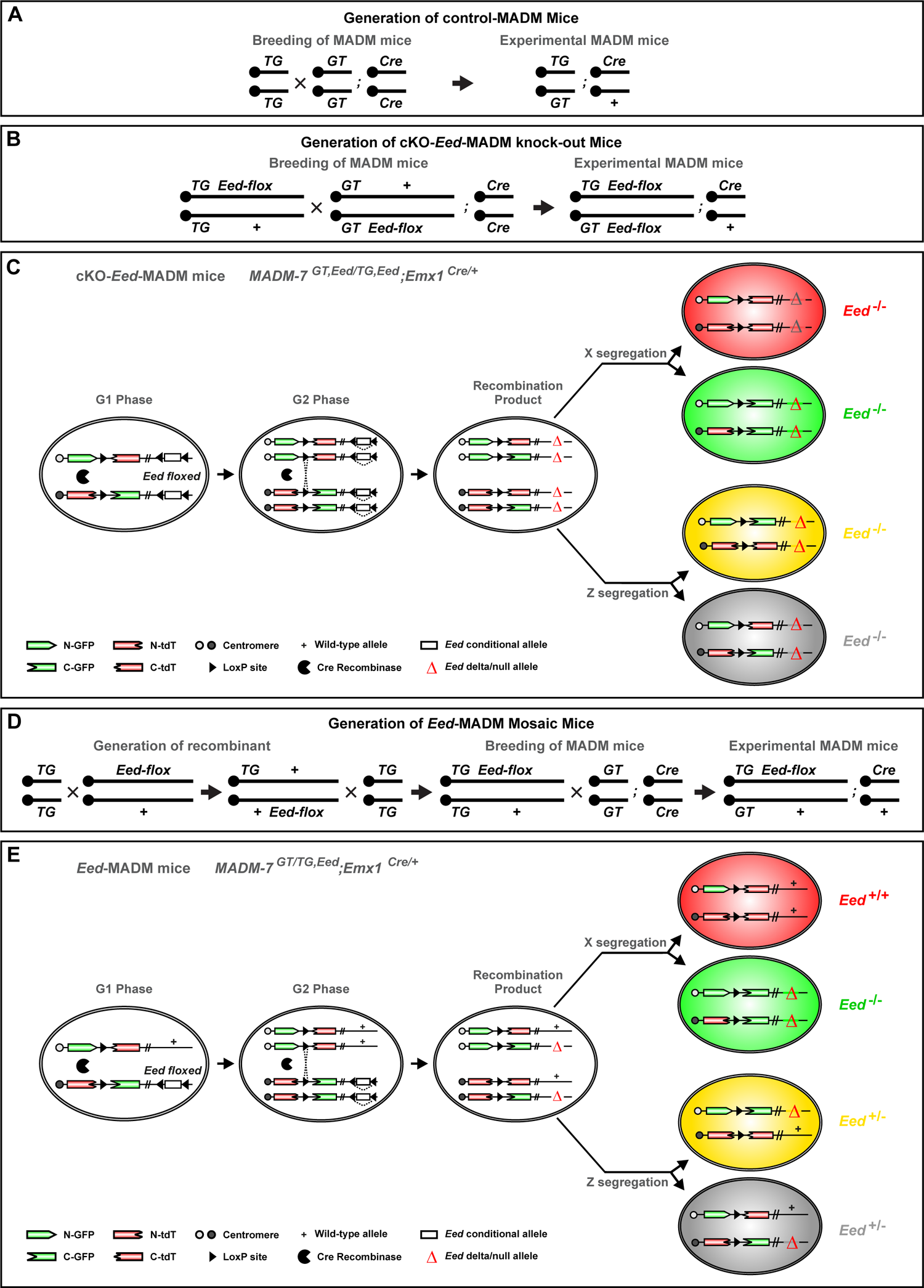
Related to Figure 1. MADM-based experimental paradigms for the generation of *Eed* mutant cells with single cell resolution. **(A)** Breeding strategy for the generation of control-MADM mice. **(B)** Breeding strategy for the generation of cKO-*Eed*-MADM mice. **(C)** MADM scheme for *Eed* conditional knockout (cKO). The *Eed* conditional allele was introduced distal to both, the GT-MADM and the TG-MADM cassettes, via meiotic recombination (Amberg and Hippenmeyer, 2021; Contreras et al., 2021). Following Cre-mediated interchromosomal recombination and mitosis, sparsely labelled GFP^+^ (green), tdT^+^ (red) and GFP^+^ tdT^+^ (yellow) homozygous *Eed* mutant cells will be generated in an unlabeled homozygous *Eed* mutant environment. **(D)** Breeding strategy for the generation of mosaic *Eed*-MADM mice. **(E)** MADM scheme for sparse mosaic *Eed* knockout. The *Eed* conditional allele was introduced distal to the TG-MADM cassette via meiotic recombination (Amberg and Hippenmeyer, 2021; Contreras et al., 2021). Upon G2-X event one GFP^+^ homozygous *Eed* mutant cell and one tdT^+^ homozygous wild-type cell will be generated in an unlabeled heterozygous environment. G2-Z segregation results in one unlabeled and one yellow heterozygous cell (Amberg and Hippenmeyer, 2021; Contreras et al., 2021). Parts of the figure were reused and adapted with permission from: (Beattie et al., 2017).

**Figure S2.**
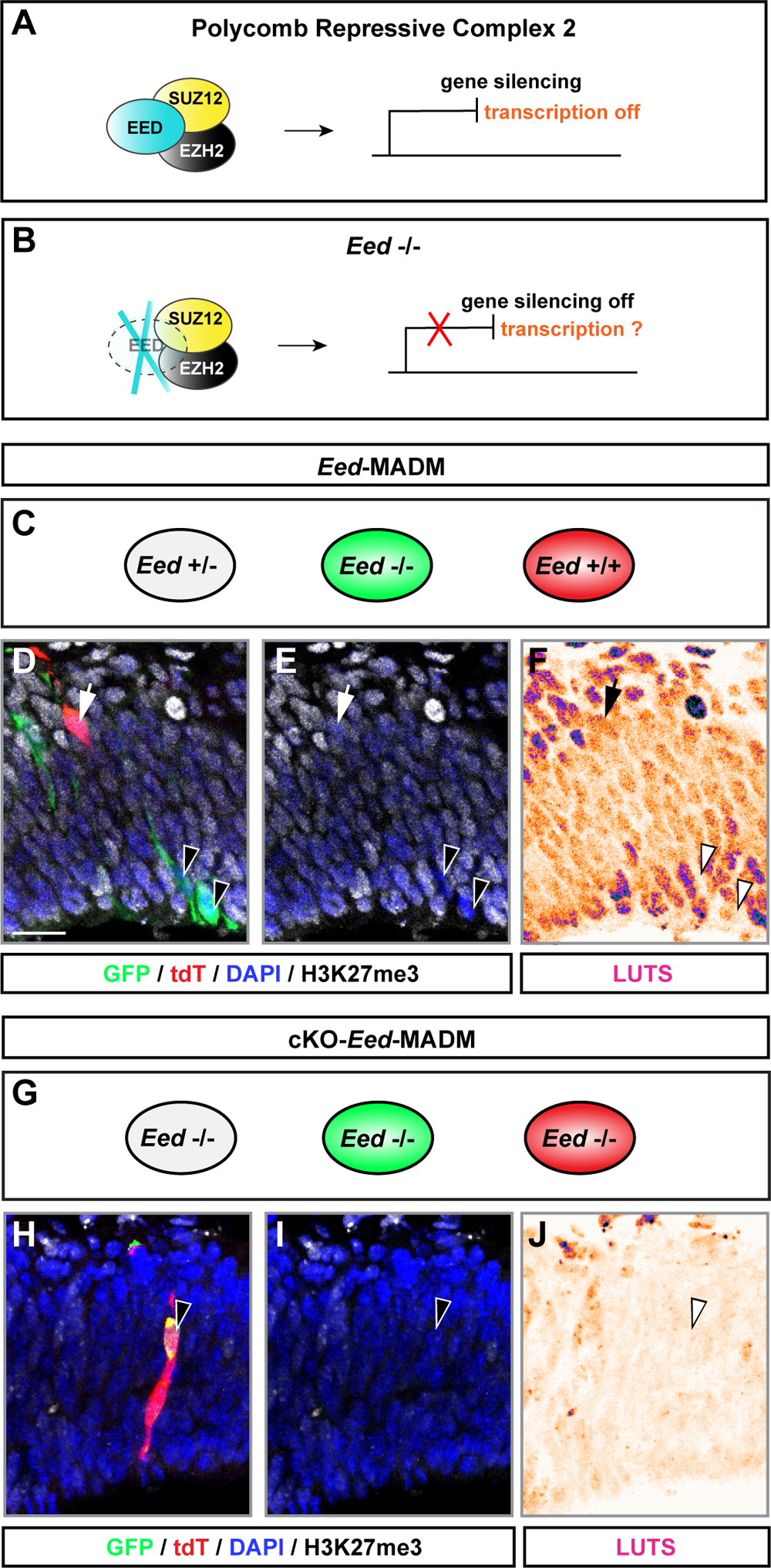
Related to Figure 1. Histological validation of PRC2 inactivation in experimental MADM paradigms. **(A)** Schematic overview of PRC2 core components and function. **(B)** Schematic overview of genetic deletion of PRC2 activity by using floxed alleles of *Eed*. **(C)** Depiction of cellular genotypes in *Eed*-MADM (sparse *Eed* deletion). **(D-F)** H3K27me3 staining in *Eed*-MADM cortex at E12.5, confirming absence of H3K27me3 and thus PRC2 activity specifically in green *Eed*^-/-^ cells. (E) H3K27me3 and DAPI only. (F) LUTS display of H3K27me3 staining intensity. Arrowheads point on individual green *Eed*^-/-^ cells which show absence of H3K27me3 mark, while full arrow points on a red *Eed*^+/+^ cell which, like the unlabeled cellular environment, is positive for H3K27me3. **(G)** Depiction of cellular genotypes in cKO-*Eed*-MADM cortex (global tissue-wide *Eed* KO). **(H-J)** H3K27me3 staining in cKO-*Eed*-MADM cortex at E12.5 confirming absence of H3K27me3 and thus PRC2 activity in all neural cells. (I) H3K27me3 and DAPI only. (J) LUTS display of H3K27me3 staining intensity. Arrowhead points on an individual yellow *Eed*^-/-^ cell which shows absence of the H3K27me3 mark, like the unlabeled cellular environment. Scale bar in (D-F, H-J): 20µm.

**Figure S3.**
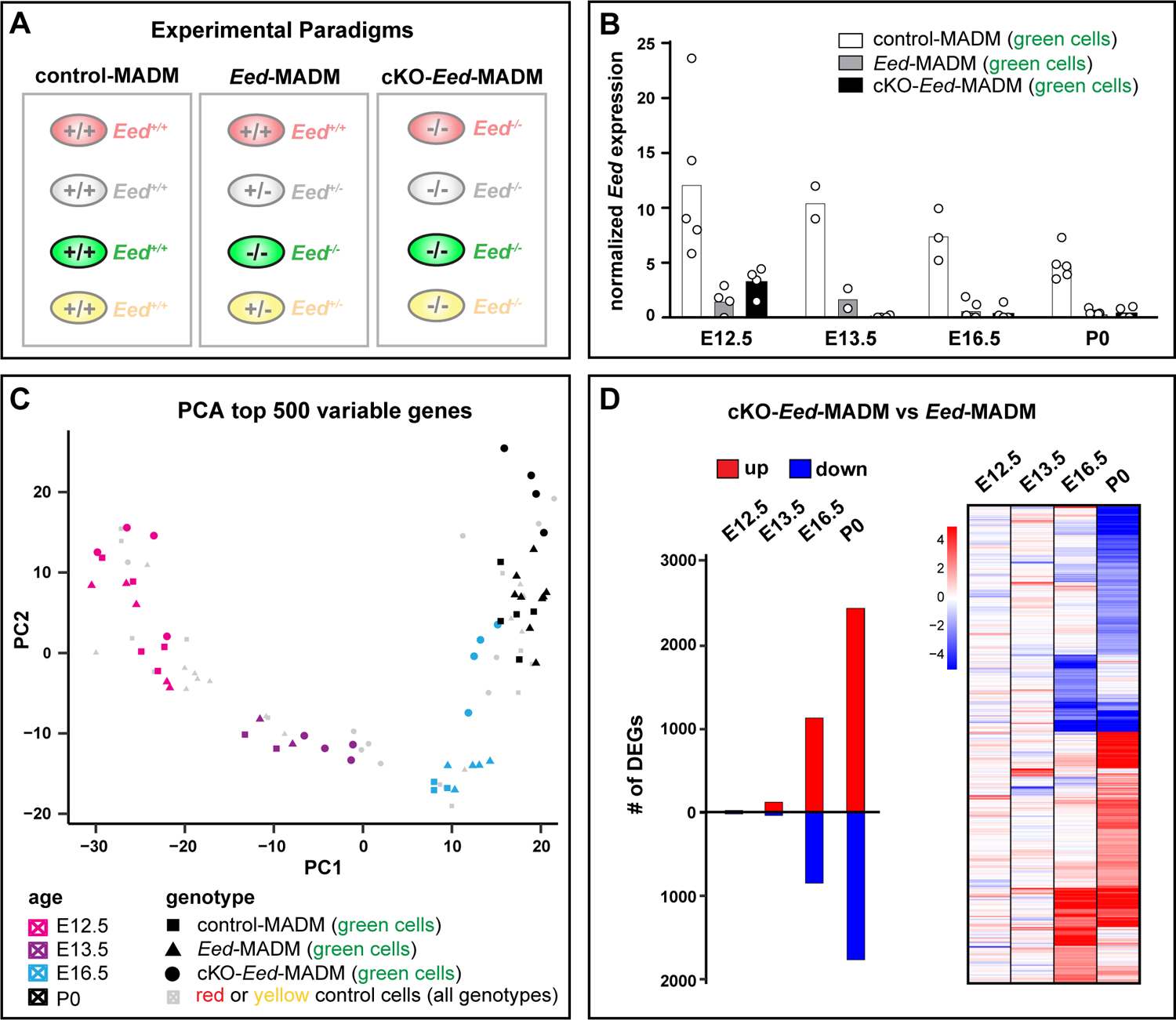
Related to Figure 3. Gene expression analysis in MADM-labeled *Eed^-/-^* cells upon sparse and global tissue-wide *Eed* deletion. **(A)** Schematic overview of experimental MADM paradigms and genotype of differentially labeled cells in control-MADM; *Eed*-MADM; and cKO-*Eed*-MADM. **(B)** Normalized *Eed* expression at E12.5, E13.5, E16.5 and P0 in purified green cells from (white) control-MADM, (grey) *Eed*-MADM and (black) cKO-*Eed*-MADM. **(C)** PCA plot of samples isolated at E12.5, E13.5, E16.5 and P0. Red and yellow control cells sorted at each time point are shown in grey. MADM-labelled green cells are color-coded according to isolation time point and displayed in pink (E12.5), purple (E13.5), blue (E16.5) and black (P0). The genetic origin of the cells is indicated by distinct geometric symbols for *Eed^+/+^* cells derived from control-MADM mice (squares), *Eed^-/-^* cells derived from *Eed*-MADM mice (triangles) and *Eed^-/-^* cells derived from cKO-*Eed*-MADM mice (circles). **(D)** Graphical representation of the number of DEGs upon comparison of *Eed^-/-^* mutant cells in *Eed*-MADM and cKO-*Eed*-MADM at 12.5, E13.5, E16.5 and P0 (left). Score heat map of DEGs from 12.5, E13.5, E16.5 and P0 upon comparison of *Eed^-/-^* mutant cells in *Eed*-MADM and cKO-*Eed*-MADM (right).

**Figure S4.**
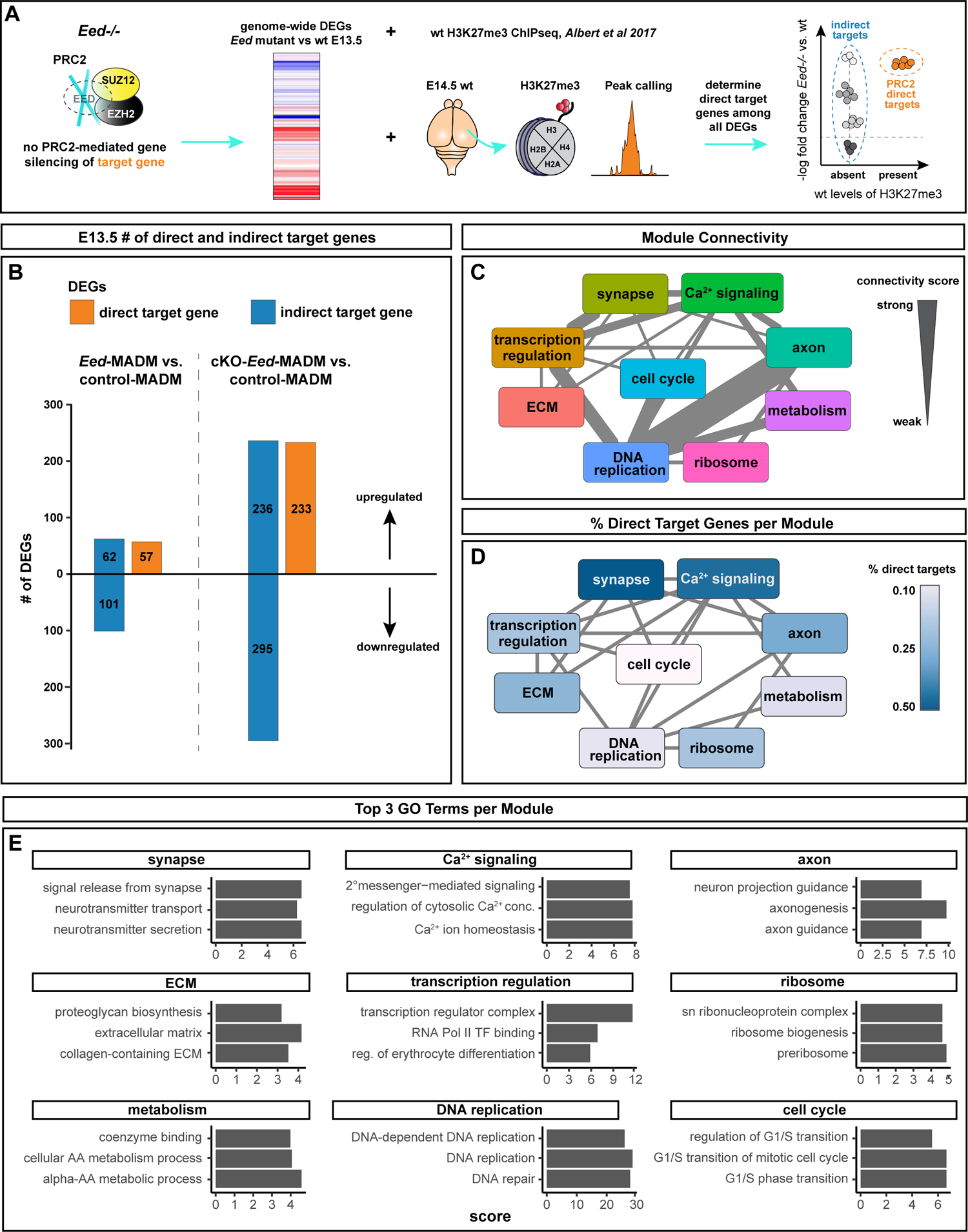
See also Figure 3. Identification and STRING network analysis of direct and indirect deregulated PRC2 target genes in *Eed^-/-^* cells upon sparse and global *Eed* deletion. **(A)** Strategy to determine direct PRC2 target genes using our RNAseq datasets from *Eed^-/-^* mutant cells aligned with H3K27me3 ChIPseq data from (Albert et al., 2017). **(B)** Bar graph showing the number of differentially expressed direct and indirect PRC2 target genes in *Eed*-MADM and cKO-*Eed*-MADM. Direct targets are indicated in orange and indirect targets in blue color. DEGs are from comparison of E13.5 *Eed^-/-^* cells isolated from *Eed*-MADM versus control-MADM and *Eed^-/-^* cells isolated from cKO-*Eed*-MADM versus control-MADM. **(C)** Top 3 Gene Ontology terms per STRING network module. **(D)** Connectivity Score of STRING network. **(E)** Bar graph showing the percentage of direct target genes per individual module in the STRING network.

**Figure S5.**
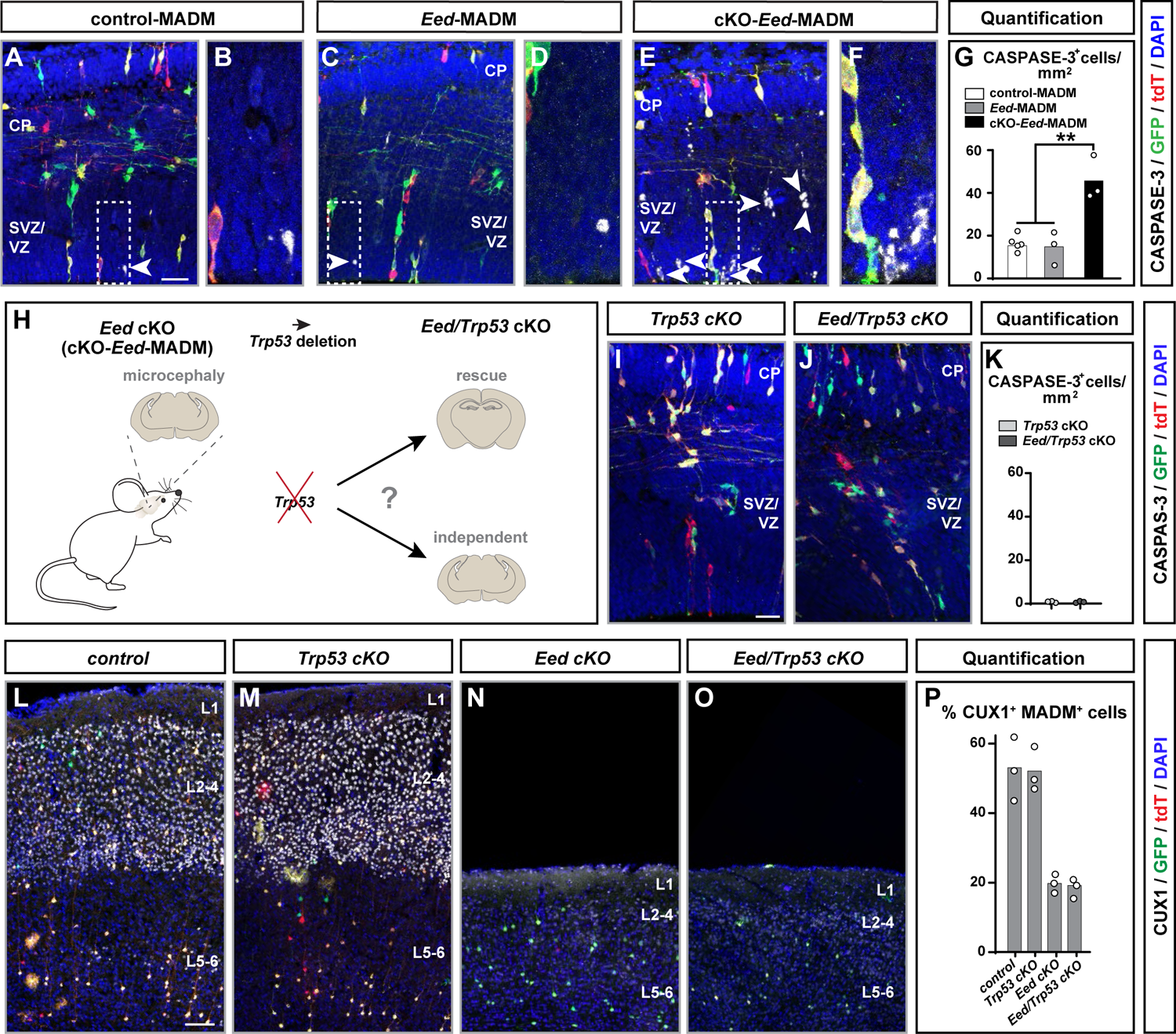
Related to Figure 4. Microcephaly in cKO-*Eed*-MADM develops in *Trp53*-independent manner. **(A-F)** Histological stainings of CASPASE-3 in (A-B) control-MADM; (C-D) *Eed*-MADM; and (E-F) cKO-*Eed*-MADM at E14.5. (B, D, F) Higher magnification of boxed areas in (A), (C) and (E) highlighting the VZ/SVZ with apoptotic cells. **(G)** Percentage of CASPASE-3^+^ cells/mm^2^ in (white) control-MADM; (grey) *Eed*-; and (black) cKO-*Eed*-MADM. **(H)** Experimental paradigm to test the possible role of *Trp53* in the emergence of microcephaly phenotype upon global KO of *Eed* in cKO-*Eed*-MADM. **(I-J)** Histological stainings of CASPASE-3 in (I) *Trp53* cKO and (J) *Eed*/*Trp53* double cKO at E14.5. **(K)** Number of CASPASE-3^+^ cells/mm^2^ in *Trp53* cKO (light grey) and *Eed*/*Trp53* double cKO (dark grey) at E14.5. **(L-O)** Confocal images with layer indications depicting immunofluorescence stainings for upper layer marker CUX1 in brains from (L) control; (M) *Trp53* cKO; (N) *Eed* cKO; and (O) *Eed/Trp53* double cKO mice at P21. **(P)** Percentage of CUX1^+^ MADM-labelled cells in brains from control, *Trp53* cKO; *Eed* cKO and *Eed/Trp53* double cKO mice at P21. Each individual data point in (G), (K) and (P) represents one experimental animal. Data indicate mean ± SEM. Statistics: (G) one-way ANOVA with multiple comparisons; * p<0.05; ** p<0.01; *** p<0.001. Data indicate mean ± SEM. Scale bars: 25µm in (A, C, E); 8µm in (B, D, F); 20µm in (I, J) and 100µm in (L-O).

**Figure S6.**
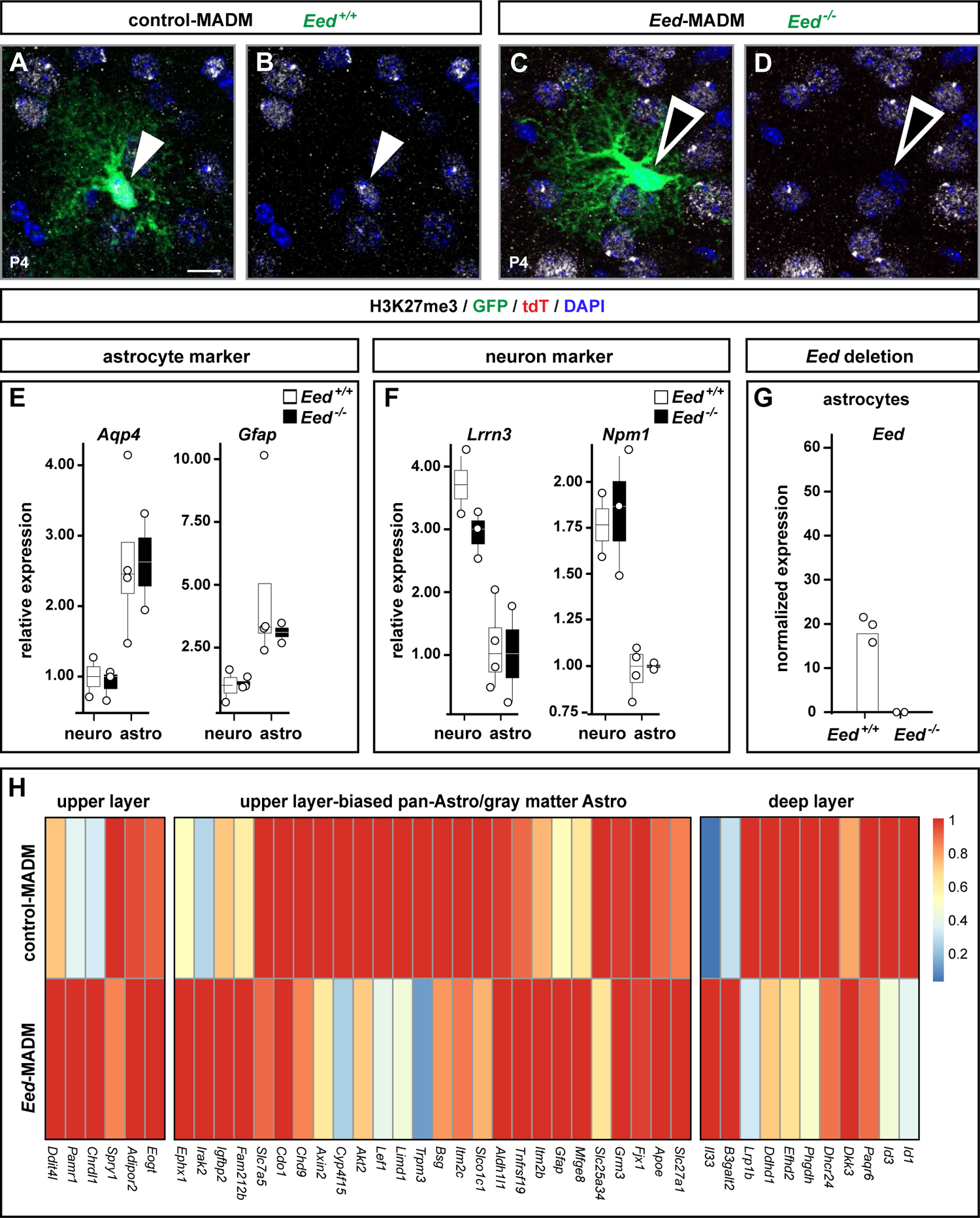
Related to Figures 5 and 6. Expression of PRC2 activity and gene expression upon *Eed* ablation in cortical astrocytes. **(A-D)** High resolution images of green GFP^+^ astrocytes in P4 (A-B) control-MADM and (C-D) *Eed*-MADM stained for H3K27me3. (B and D) Images from (A and C) showing H3K27me3 and DAPI only. Closed white arrowheads point on an immature *Eed^+/+^* astrocyte with H3K27me3 marks. Black arrowheads with white lining point on an immature *Eed^-/-^* astrocyte devoid of the H3K27me3 marks. **(E-F)** Assessment of purity of P4 GFP^+^/lacZ^+^ (astrocyte) and GFP^+^/lacZ^-^ (neuron) cell populations by comparing the expression of (E) astrocyte marker genes *Aqp4* and *Gfap* and (F) neuron marker genes *Lrrn2* and *Npm1* in purified *Eed^+/+^* (white) and *Eed^-/-^* (black) neuron and astrocyte cell populations. Abbreviations: neuro – neurons; astro – astrocytes. **(G)** Normalized *Eed* expression in purified *Eed^+/+^* (white) and *Eed^-/-^* (black) astrocytes isolated from control-MADM and *Eed*-MADM at P4. **(H)** Expression of cortical astrocyte marker genes as defined in (Bayraktar et al., 2020) from *Eed^+/+^* and *Eed^-/-^* astrocytes isolated at P4. Each individual data point represents one experimental animal. Data show mean ± SEM. Scale bar in (A-D): 10µm.

## Supplementary Table Legends

**Table S1. Complete results of DEG, H3K27me3 and GO term enrichment analysis.** Tabs labeled with DEG contain results from DESeq2 analyses, tabs labeled with GO contain results of GO term enrichment analyses from clusterProfiler. Tab DEG.E13_cKO_wt contains additional information on STRING analysis (cluster, pred_function) and H3K27me3 analysis (column direct_H3K27me3). Column cluster indicates arbitrary numbering of clusters in the STRING network and are the same as for the labels of the GO term enrichment analyses. Abbreviations: cKO: cKO-*Eed*-MADM, het: *Eed*-MADM, wt: control-MADM

**Table S2.** Complete results of astrocyte DEG analysis. Table contains output from astrocyte DEG analysis using DESeq2.

**Table S3.** Statistics table. Table contains the number of mice, sections, quantified cells and parameters for all figures.

